# Robust, data-driven bioregionalizations emerge from diversity concordance

**DOI:** 10.1101/2021.08.31.458457

**Authors:** Cristian S. Montalvo-Mancheno, Jessie C. Buettel, Stefania Ondei, Barry W. Brook

## Abstract

**Aim:** Despite the increasing interest in developing new bioregionalizations and assessing the most widely accepted biogeographic frameworks, no study to date has sought to systematically define a system of small bioregions nested within larger ones that better reflect the distribution and patterns of biodiversity. Here, we examine how an algorithmic, data-driven model of diversity patterns can lead to an ecologically interpretable hierarchy of bioregions.

**Location:** Australia.

**Time period:** Present.

**Major taxa studied:** Terrestrial vertebrates and vascular plants.

**Methods:** We compiled information on the biophysical characteristics and species occupancy of Australia’s geographic conservation units (bioregions). Then, using cluster analysis to identify groupings of bioregions representing optimal discrete-species areas, we evaluated what a hierarchical bioregionalization system would look like when based empirically on the within-and between-site diversity patterns across taxa. Within an information-analytical framework, we then assessed the degree to which the World Wildlife Fund’s (WWF) biomes and ecoregions and our suite of discrete-species areas are spatially associated and compared those results among bioregionalization scenarios.

**Results:** Information on biodiversity patterns captured was moderate for WWF’s biomes (50– 58% for birds’ beta, and plants’ alpha and beta diversity, of optimal discrete areas, respectively) and ecoregions (additional 4–25%). Our plants and vertebrate optimal areas retained more information on alpha and beta diversity across taxa, with the two algorithmically derived biogeographic scenarios sharing 86.5% of their within- and between-site diversity information. Notably, discrete-species areas for beta diversity were parsimonious with respect to those for alpha diversity.

**Main conclusions:** Nested systems of bioregions must systematically account for the variation of species diversity across taxa if biodiversity research and conservation action are to be most effective across multiple spatial or temporal planning scales. By demonstrating an algorithmic rather than subjective method for defining bioregionalizations using species-diversity concordances, which reliably reflects the distributional patterns of multiple taxa, this work offers a valuable new tool for systematic conservation planning.

## 1. Introduction

The division of the Earth’s surface into regions of unique biotic communities or similar ecological processes is a cornerstone of biogeographic and macroecological research (Ebach & Parenti, 2015; Mackey, Berry, & Brown, 2008). The identification of alternative macrounits of biodiversity reflects differences in scale of analysis, methods of classification, and type of data governing the existing biogeographic frameworks (Mackey et al., 2008). Many bioregionalizations have delineated precise geographic units based on differences in species composition (Kreft & Jetz, 2010; Wallace, 1876) and/or discontinuities in the abiotic environment (Olson et al., 2001; Omernik, 2004). Recently, biogeographic frameworks have focused instead on either using phylogenetic data (Daru, Elliott, Park, & Davies, 2017; Holt et al., 2013; Maestri & Duarte, 2020) to define those hard boundaries, or have taken a ‘softer’ approach to their geographic delineation by identifying transition zones (Edler, Guedes, Zizka, Rosvall, & Antonelli, 2017; Vilhena & Antonelli, 2015). Another key difference is that while some biogeographers have sought to define systems of small units nested within larger ones— such is the case of World Wildlife Fund (WWF) Terrestrial Ecoregions (Dinerstein et al., 2017; Olson et al., 2001)—others have rejected this as a desirable outcome of the study system (Ebach & Parenti, 2015). At finer resolution, the spatial delineation of homogeneous areas characterized by broad, landscape-scale natural features and environmental processes (called land systems, bioregions, or equivalents) has also been crucial in the development of many bioregionalizations at sub-continental scales (Mackey et al., 2008). Yet, despite the great contribution that this diversity of frameworks has made to our understanding of biodiversty patterns, creating an objective, repeatable and transferable hierarchical system of geographic operational units to meaningfully aggregate biodiversity from-regional-to-global scale, across multiple taxonomic groups, remains elusive (Antonelli, 2017; Morrone, 2018).

Alongside Wallace’s zoogeographic regions (Wallace, 1876), WWF’s hierarchical framework (Olson et al., 2001) of discrete areas of natural communities (i.e., ecoregions), spatially nested within larger distinct areas that reflect relations between climate, flora and fauna (i.e., biomes), is the most widely accepted global bioregionalization that has been the foundation of scientific research, environmental policy, resource management and conservation for almost two decades (Kier et al., 2005; Mackey et al., 2008; Smith et al., 2018). However, few continental-level bioregionalizations that correspond to a spatial subdivision of WWF’s geographic operational units have been developed and adopted over this period (Omernik, 2004; Thackway & Cresswell, 1995). Notably, only the bioregionalization for Australia—known as the Interim Biogeographic Regionalization for Australia (IBRA) framework—has been explicitly defined as a more detailed geographic division of WWF’s ecoregions (Department of Agriculture, 2012a).

Since its inception, the IBRA framework has been framed as a tool to guide the systematic conservation planning of Australia’s biodiversity. The initial release divided the continent in 80 biogeographic regions, called bioregions (Thackway & Cresswell, 1995). Over the subsequent 25 years, new bioregions have been identified, and boundaries updated to coincide better with faunal and floral patterns and environmental processes that influence the functioning of entire ecosystems. In an effort to aid regional conservation, IBRA bioregions have been further divided into subregions, using finer-scale differences in biophysical attributes— such as geology and vegetation—to spatially define major regional ecosystems. Like WWF’s global ecoregions and other continental-scale bioregionalization templates (Mackey et al., 2008; Omernik, 2004), the IBRA framework reflects a hierarchical structure delineated bio-topically where the spatial aggregation of subregions makes up bioregions (Department of Agriculture, 2012a). The creation and update of WWF’s and Australia’s biogeographic frameworks have also been similar, in that the tacit knowledge of an expert panel was used to compile a suite of disparate spatial information to define regions within which geographic phenomena associated with differences in ecosystems’ characteristics (i.e., health, quality, and integrity) coincide (Dinerstein et al., 2017; Olson et al., 2001; Omernik, 2004; Thackway & Cresswell, 1995). The subjective, expert-based derivation of these two bioregionalizations has prompted criticism. Nonetheless, only WWF’s biomes and ecoregions have been scrutinized quantitatively, with their capacity to discriminate species diversity shown to perform better than a random allocation of boundaries (Smith et al., 2018), but worse in comparison to remotely sensed productivity clusters (Coops, Kearney, Bolton, & Radeloff, 2018).

A plethora of studies seeking to develop new bioregionalization scenarios, and to assess the most widely accepted biogeographic templates, have emerged over the last decade (Ebach & Parenti, 2015; Kreft & Jetz, 2010). This revived interest is due to a number of recent developments. There is an increasing accessibility to ecological datasets that provide systematic information on species distributions, as well as on other facets of biodiversity (e.g., phylogenetic, and functional diversity) over large extents and at an increasingly finer resolution for many taxa (Daru et al., 2017; Ficetola, Mazel, & Thuiller, 2017). Alongside this, ecologists and biogeographers are increasingly using remotely sensed data to understand biodiversity patterns and process across multiple spatial and/or temporal scales (Coops et al., 2018). Nonetheless, many—if not all—of the aggregative frameworks to emerge during the past 20 years would not have been possible without the use of high-performance computing infrastructure. Together with high volumes of processing power, spatially explicit aggregative and comparative techniques and new approaches to disentangle fundamental properties of ecological systems have also been introduced. These advances have opened the possibility to develop new algorithm-driven bioregionalizations that are objective, reproduceable, and tractable. However, whether the operational units of quantitative bioregionalizations—like in bioregionalizations defined by an expert-panel—can capture multiple facets of biodiversity remains highly contested and of much research interest among biogeographers (Ebach & Parenti, 2015; Mackey et al., 2008; Morrone, 2018).

Motivated by these problems with definition and implementation, we developed an integrative, data-driven approach to bioregionalization that leverages the information on species richness and composition within the bioregions of the IBRA framework. More specifically, we asked: 1) What would an IBRA framework look like when based empirically on the accumulation of species with increase in area (species-area relationship), and the within- and between-site species diversity (alpha and beta) for multiple taxonomic groups (hereafter referred as optimal discrete-species clusters)? 2) How well do WWF’ biomes and ecoregions match ‘optimal’ discrete-species clusters? 3) Is the spatial configuration of optimal-discrete-plant and - vertebrate clusters better associated with each other than with other discrete species clusters at lower taxonomic ranks, and when compared to the spatial association between WWF’s operational units (biomes and ecoregions) and those of optimal-discrete species clusters? By answering those questions, we reveal a hierarchical system of spatial partitions that is ecologically interpretable, and thereby best suited to inform biodiversity policy, research, and conservation.

## 2. Materials and Methods

### 2.1 Data collection and processing

We collected spatial information on climate (Hallgren et al., 2016), elevation (Earth Resources Observation and Science (EROS) Center, NA), vegetation (Department of Agriculture, 2018), soil (Australian Soil Resource Information System, 2013), lithology (Raymond et al., 2012), and occurrences of terrestrial species native to Australia (Atlas of Living Australia, NA), which after pre-processing to minimize errors and biases included 25,995 native species: 23,248 vascular plants, 233 amphibians, 1,201 birds, 349 mammals, and 964 reptiles (see Appendix S1 in Supporting Information for details).

We downloaded version seven of IBRA subregions’ names and borders (Department of Agriculture, 2012b) to derive a spatially coherent four-tier hierarchical system of geographic operational units for Australia (see Appendix S1 for details)—where 410 IBRA subregions are nested within 85 IBRA bioregions. Based on the most recent version of WWF’s bioregionalization (Dinerstein et al., 2017) those bioregions are nested within 37 ecoregions, and those macrounits are themselves embedded in 7 broader-scale (and spatially coherent) biomes. We chose to use geographic operational units, because analyses based on lists of species within such units, as opposed to a grid-based approach, can highlight gradual changes in species diversity and are less likely to distort areal relationships due to heterogeneity in the sizes of species ranges (Kreft & Jetz, 2010; Kreft, Sommer, & Barthlott, 2006; Morrone & Escalante, 2002). We deem this a desirable feature since our goal was to reveal a bioregionalization’s hierarchical system of discrete spatial clusters that is more directly relevant to biodiversity.

We characterized Australia’ biophysical space by calculating the mean value of elevation and of nineteen climatic variables within IBRA subregions and bioregions. As the geographic-based measures for lithology, soil, and vegetation, we computed the percentage cover for each of these factors’ categories relative to the size of subregions and bioregions (see Appendix S1 for details on excluded categories for these three discrete variables). Meanwhile, for the characterization of Australia’s biotic space, we derived presence-absence matrices for vascular plants and four vertebrate classes (amphibians, birds, mammals, and reptiles) by intersecting IBRA bioregions and subregions with both species occurrences with less than 20 records post-equalization (i.e., a procedure to even out the difference in number of species occurrences among IBRA subregions by minimizing the variance within subregions’ size-classes, while maximizing the variance between size classes) and our set of empirical extent-occurrence maps for those species with at least 20 records (see Appendix S1 for details and rationality). We joined amphibian and reptile presence-absence matrices into a single group (herpetofauna) to ensure that there were at least 10 species per IBRA operational units across taxonomic groups; we also created a presence-absence matrix for all vertebrate species. In terms of species, amphibians and reptiles follow different biogeographic patterns (Powney, Grenyer, Orme, Owens, & Meiri, 2010), yet as a broad taxonomic group (i.e., herpetofauna), exothermic species represent a huge array of evolutionary adaptations that allow them to cover a wide range of potential niches. We used ArcGIS v. 10.5.1 (2017) to harmonize spatial data to a common format and coordinate reference system (Australian Albers Equal Area; EPSG: 3577). All spatial calculations and feature engineering were done in R v. 3.6.3 (R Core Team, 2020) using several packages (see Appendix S1 for complete list).

### 2.2 Metric to discriminate spatial clusters in biophysical space

To assess the ecological significance of the ordination of geographic operational units of the IBRA framework based on biophysical factors, we used the species-area relationship (SAR) as a metric because SAR is one of the well-studied properties of ecological systems and has been applied in identifying priority areas for biodiversity conservation at large scales (Guilhaumon, Gimenez, Gaston, & Mouillot, 2008; Triantis, Guilhaumon, & Whittaker, 2012). Among nine alternative mathematical functions, we selected the logarithmic form of the power function to fit SAR for vascular plants and selected vertebrate species (bird, mammal, and herpetofauna) based on Akaike’s Information Criteria (AIC) (Akaike, 1974) (see Appendix S2 for details and results).

### 2.3 Selection of biophysical factors and IBRA unit of analysis

Our selected geographic and environmental covariates represent a complex dataset (n = 71 variables) that describes IBRA subregions and bioregions by sets of variables structured into groups. We defined the distance between distinct IBRA spatial clusters at a hierarchical level to be based on an equal contribution of these five groups of continuous variables. To balance the influences of each group of variables in the description of distinct spatial clusters (Bécue-Bertaut & Pagès, 2008), we used multiple factor analysis (MFA) to assess the contribution of groups of variables to the characterization of IBRA operational units, and to identify the number of principal components needed to retain at least 90% of the variance, using their eigenvalues to model the dissimilarity of IBRA operational units in the biophysical space instead of using geographic-based measures.

Additionally, we assessed the relevance of MFA results to discriminate biophysical factors, if necessary, and to identify the most appropriate hierarchical level of the IBRA framework for revealing the nature of the IBRA framework differences in terms of species diversity. We did this by visually exploring the spatial coherence and ecological interpretability of the ordination of subregions and bioregions into seven clusters—matching the number of WWF biomes—based on principal components and the ‘static’ technique to cut dendrograms (see next section for details). This assessment identified IBRA bioregions as the most appropriate geographic unit of analysis compared with subregions and reduced the biophysical dataset’s structure to include the eigenvalues of the 27 principal components based on climate, lithology, soil, and vegetation (see Appendix S3 for details) when constructing the ordination structure of IBRA bioregions in biophysical space. We used ‘FactoMineR’ v. 1.42 (Lê, Josse, & Husson, 2008) package in the program R to perform MFA, with variables standardized, and the name of IBRA subregions and bioregions set as non-active variables.

### 2.4 Discontinuities of species diversity

#### 2.4.1 Hierarchical clustering

We ordered IBRA bioregions using the Ward’s method as the clustering algorithm, and Euclidian distance and species turnover—measured with the Beta-Simpson index (Lennon, Koleff, GreenwooD, & Gaston, 2001; Simpson, 1943)—as the dissimilarity measures of bioregions in terms of principal components for our suite of biophysical factors (hereafter referred as PC-biophysical) and species composition, respectively. Since no existing method is capable of maximizing both clustering criteria simultaneously (i.e., the amount of information retained in the dendrogram, and the clusters’ internal coherence), we chose to use Ward’s algorithm, because it has proven to perform best in the second criterion (Castro-Insua, Gómez-Rodríguez, & Baselga, 2018; Kreft & Jetz, 2010). In addition, the identification of a hierarchical system of distinct spatial clusters that minimizes within-cluster and maximizes between-cluster dissimilarity (i.e., clusters’ internal coherence) in terms of biodiversity is a highly desirable outcome for any bioregionalization (Ebach & Parenti, 2015; Kreft & Jetz, 2010), and aligns clearly with our study’s overarching goal. We used the ‘stats’ v. 3.6.3 (R Core Team, 2020), and the ‘betapart’ v. 1.5.1 (Baselga & Orme, 2012) packages to compute dissimilarity matrices. While bioregions’ cluster analysis based on species compositional dissimilarity was done using ‘stats’ too, we used the ‘FactoMineR’ (Lê et al., 2008) package to conduct that analysis in the PC-biophysical dissimilarity space.

#### 2.4.2 Optimal discrete-species clusters

We cut the dendrograms resulting from the hierarchical cluster analyses of IBRA bioregions in the PC-biophysical space and the dissimilarity in species composition for five taxonomic groups (birds, mammals, herpetofauna, vertebrate, and vascular plants) using different techniques to identify sets of prominent, spatially coherent biodiversity clusters. This included: defining continuous dendrogram branches based on a desired number of clusters (‘static’ technique), pruning branches based on their structure in the dendrogram (‘dynamic’ technique; specifically a bottom-up algorithm, called Dynamic Hybrid Cut) (Langfelder, Zhang, & Horvath, 2008), and/or identifying the intersection point between two straight lines that best fit an evaluation curve by minimizing the total root mean square error (‘L’ technique) (Salvador & Chan, 2004). We used these pruning techniques together with goodness-of-fit and parsimony metrics (e.g., R-squared, and AIC and/or Bayesian Information Criteria – BIC) to discriminate prominent clusters in terms of species accumulation as the area sampled is increased (SAR models for prominent clusters in the PC-biophysical space), and the within- and across-variance of species richness and composition (alpha and beta diversity models, respectively). We selected the most parsimonious yet ecologically coherent model of the species-area relationship, and pairs of the best models of the variance in alpha and beta diversity, respectively, as the optimal discrete-species clusters (see Appendix S4 for methodological details).

### 2.5 An ecologically meaningful algorithmic IBRA framework

To reveal a hierarchical system of IBRA bioregions that meaningfully aggregates species diversity from-regional-to-global scale, we examined the ecological interpretability of our suite of eleven optimal discrete-species clusters (i.e., ten optimal partitions of species diversity, along with the optimal partition of SAR). We did this by calculating the slope and the standard error of the regression line of the species-area relationship across five taxonomic groups for (i) the spatial configuration of the optimal SAR model and (ii) the bespoke WWF’s classification of IBRA bioregions into seven biomes. We then plotted the distribution, mean, median, and standard deviation of the merging height of nodes in the dendrograms of species composition, to visualize differences in the dissimilarity distance within and across the five target taxonomic groups.

We assessed the spatial coherence of optimal discrete-species clusters visually, and then quantitatively by estimating how well WWF’s classification of IBRA bioregions into biomes and ecoregions captured information on species richness and turnover stored in our set of maps for the optimal discrete-species clusters. We computed an overall global measure of association— called the ‘V-measure’ and implemented in the ‘sabre’ v. 0.3.2 (Nowosad & Stepinski, 2018) package—to quantify the degree of spatial association between these maps. The 0-1 range of the V-measure is grounded in information theory and interpretable in terms of analysis of variance, where 0 indicates absence of spatial association between two maps and 1 when the spatial association is perfect–respectively meaning that the amount of mutual information of a pairwise-map comparison totally differs or is identical. We computed this global measure of association to determine whether the amount of information on species richness and composition stored in the optimal discrete-plant clusters and the optimal discrete-vertebrate clusters is higher with each other than with other discrete-species clusters at lower taxonomic ranks, and then compared these results with those for the WWF ecoregions.

## 3. Results

### 3.1 Optimal discrete-species clusters

To identify an optimal partition of IBRA bioregions in the PC-biophysical space, we fitted 150 SAR models across five taxonomic groups to five sets of prominent clusters that were defined using three different dendrogram cutting techniques (see methods and Appendix S4 for details). Based on AIC scores, the best-selected SAR models across this set of prominent clusters frequently included the same grouping of bioregions as the best distinct spatial cluster among five taxonomic groups, except in three cases: SAR of plants where the seven groups were defined using the ‘dynamic’ technique, and SAR of mammals in prominent clusters with seven and nine groups based on ‘static’ and the ‘L’ techniques, respectively (Table 1).

**Table 1.**
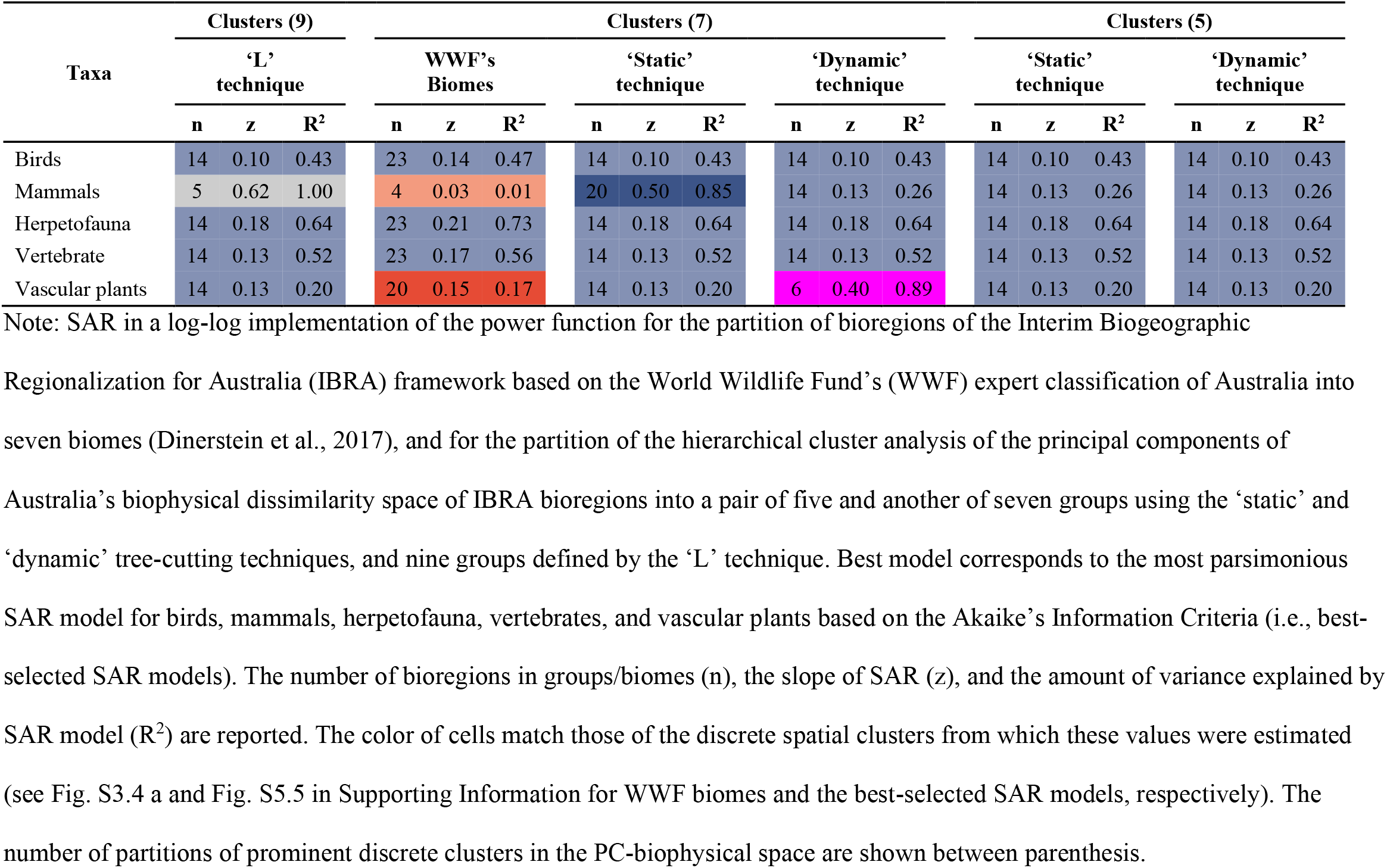
Best models of the species-area relationship (SAR).

| Taxa | Clusters (9) |  |  | Clusters (7) |  |  |  |  |  |  |  |  | Clusters (5) |  |  |  |  |  |
| --- | --- | --- | --- | --- | --- | --- | --- | --- | --- | --- | --- | --- | --- | --- | --- | --- | --- | --- |
|  | ‘L’ technique |  |  | WWF’s Biomes |  |  | ‘Static’ technique |  |  | ‘Dynamic’ technique |  |  | ‘Static’ technique |  |  | ‘Dynamic’ technique |  |  |
|  | n | z | R<sup>2</sup> | n | z | R<sup>2</sup> | n | z | R<sup>2</sup> | n | z | R<sup>2</sup> | n | z | R<sup>2</sup> | n | z | R<sup>2</sup> |
| Birds | 14 | 0.10 | 0.43 | 23 | 0.14 | 0.47 | 14 | 0.10 | 0.43 | 14 | 0.10 | 0.43 | 14 | 0.10 | 0.43 | 14 | 0.10 | 0.43 |
| Mammals | 5 | 0.62 | 1.00 | 4 | 0.03 | 0.01 | 20 | 0.50 | 0.85 | 14 | 0.13 | 0.26 | 14 | 0.13 | 0.26 | 14 | 0.13 | 0.26 |
| Herpetofauna | 14 | 0.18 | 0.64 | 23 | 0.21 | 0.73 | 14 | 0.18 | 0.64 | 14 | 0.18 | 0.64 | 14 | 0.18 | 0.64 | 14 | 0.18 | 0.64 |
| Vertebrate | 14 | 0.13 | 0.52 | 23 | 0.17 | 0.56 | 14 | 0.13 | 0.52 | 14 | 0.13 | 0.52 | 14 | 0.13 | 0.52 | 14 | 0.13 | 0.52 |
| Vascular plants | 14 | 0.13 | 0.20 | 20 | 0.15 | 0.17 | 14 | 0.13 | 0.20 | 6 | 0.40 | 0.89 | 14 | 0.13 | 0.20 | 14 | 0.13 | 0.20 |

Note: SAR in a log-log implementation of the power function for the partition of bioregions of the Interim Biogeographic Regionalization for Australia (IBRA) framework based on the World Wildlife Fund's (WWF) expert classification of Australia into seven biomes (Dinerstein et al., 2017), and for the partition of the hierarchical cluster analysis of the principal components of Australia's biophysical dissimilarity space of IBRA bioregions into a pair of five and another of seven groups using the 'static' and 'dynamic' tree-cutting techniques, and nine groups defined by the 'L' technique. Best model corresponds to the most parsimonious SAR model for birds, mammals, herpetofauna, vertebrates, and vascular plants based on the Akaike's Information Criteria (i.e., best- selected SAR models). The number of bioregions in groups/biomes (n), the slope of SAR (z), and the amount of variance explained by SAR model (R<sup>2</sup>) are reported. The color of cells match those of the discrete spatial clusters from which these values were estimated (see Fig. S3.4 a and Fig. S5.5 in Supporting Information for WWF biomes and the best-selected SAR models, respectively). The number of partitions of prominent discrete clusters in the PC-biophysical space are shown between parenthesis.

These three special cases also had the highest variance in species richness explained by size of bioregions (Table 1). In the prominent clusters to which two of them belong (i.e., prominent clusters with seven and nine groups based on ‘static’ and the ‘L’ techniques), the Tropical and Subtropical Moist Broadleaf Forests biome was reconstructed (Fig. S5.5 c and e in Supporting Information). Yet, the rate of increase in the number of species per standard area differed greatly among taxonomic groups for this biome (Biome 7 in Fig. 1b), with the slope of SAR models for vertebrates and vascular plants being 1.5 and 5.5 the slope of the bird SAR model. Considering this as a whole, the partition of bioregions into seven groups using the ‘dynamic’ tree-cutting technique—which based on its formulation is a technique that improves the detection of outlying group members in a prominent cluster (Langfelder et al., 2008)—was finally selected as the optimal partition for changes in species richness per standard area (i.e., optimal partition for SAR).

**FIGURE 1.**
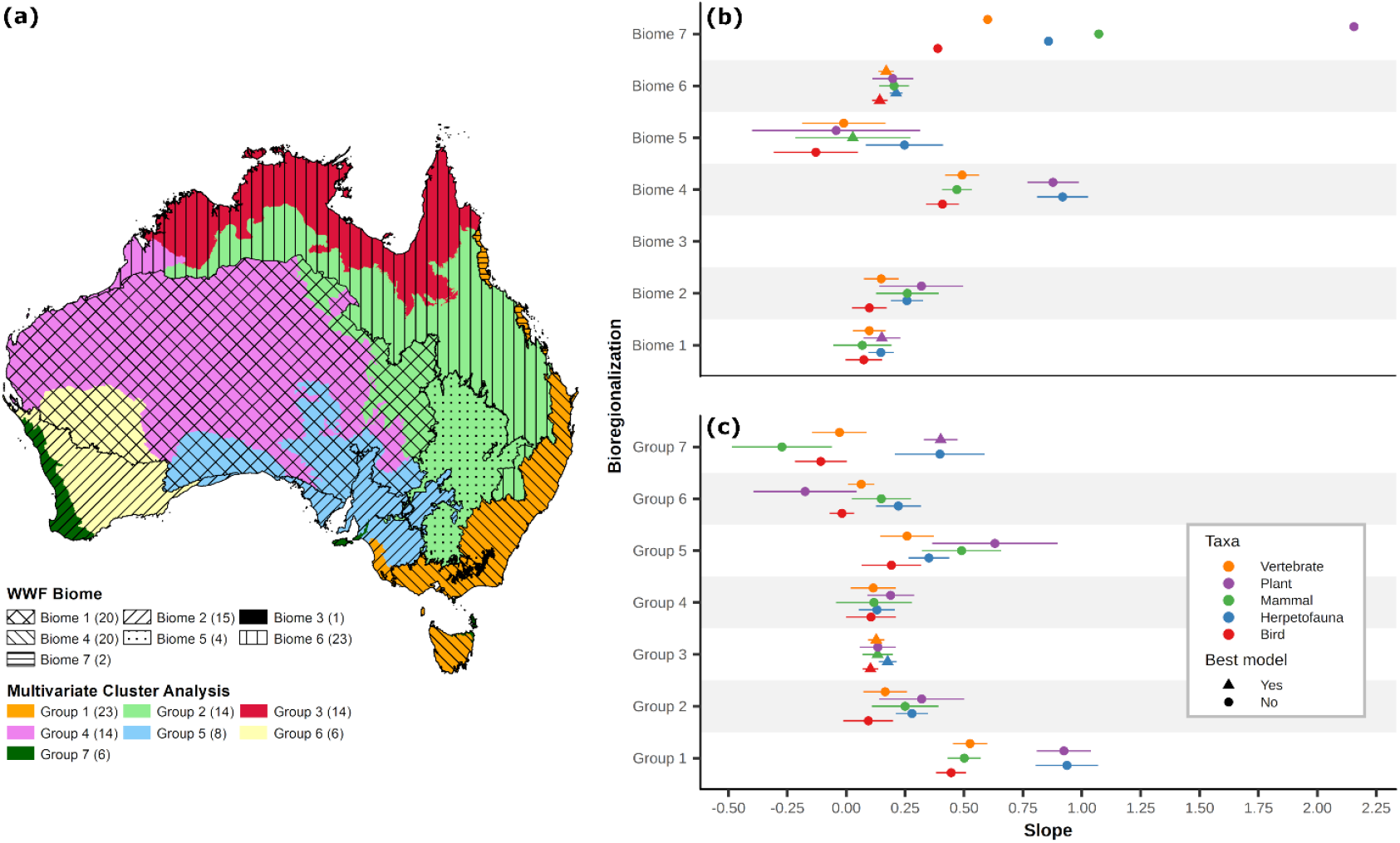
Relationship between biophysical factors and species richness. (a) Overlap of distinct spatial clusters of Australia’s bioregions based on the World Wildlife Fund’s (WWF) biomes (Dinerstein et al., 2017), and the optimum partition for changes in species richness per standard area (i.e., optimal partition for SAR), with the number of bioregions in biomes/groups found between parentheses. Species-area relationship (SAR) in a log-log implementation of the power function for the aggregations of bioregions according to (b) WWF’s biomes and (c) groups of the optimal partition for SAR. Point represents the slope (z-value) of the shape of SAR, and line shows the standard error of the regression line. Due to Biome 3 including only one bioregion, neither slope nor the standard error could be computed, whereas only the slope could be estimated for Biome 7. Biome 1 = Deserts & Xeric Shrublands; Biome 2 = Mediterranean Forests, Woodlands, Scrub; Biome 3 = Montane Grasslands & Shrublands; Biome 4 = Temperate Broadleaf & Mixed Forests; Biome 5 = Temperate Grasslands, Savannas & Shrublands; Biome 6 = Tropical & Subtropical Grasslands, Savannas & Shrublands; and Biome 7 = Tropical & Subtropical Moist Broadleaf Forests.

Based on our approach to discriminate prominent clusters in terms of the variance of alpha or beta diversity across groups of bioregions in sets of prominent clusters (see Appendix S4), the clearest break in log-likelihood of the within- and across-variance of species richness (i.e., ANOVA models for birds, mammals, herpetofauna, vertebrate, and vascular plants) was evident when dendrograms of compositional dissimilarity were cut using the ‘dynamic’ technique (Fig. S6.6). Notably, the optimal discrete-species cluster for vertebrates’ alpha diversity was defined using only the prominent cluster with the lowest BIC score rather than the agreement between AIC and BIC scores, as done for the other taxonomic groups. Likewise, the highest ratio between the sum of multiple-site measures of compositional dissimilarity across groups of bioregions in sets of prominent clusters and the multiple-site measure of compositional dissimilarity across Australia’s bioregions appeared stable across taxa—particularly for birds, mammals, and vascular plants—when prominent clusters were defined by ‘dynamic’ tree-cutting technique (Fig. S7.7). As in the optimal partition for SAR, optimal partitions of alpha and beta diversity among all five taxonomic groups were also more appropriately identified when dendrograms of species compositional dissimilarity were cut using the ‘dynamic’ technique, and thereby this suite of eleven optimal discrete-species clusters were used in subsequent analysis.

### 3.2 An ecologically meaningful IBRA framework

Using the spatial configuration of our suite of optimal clusters (Fig. S8.8 b–l) to examine what an IBRA framework would look like when based empirically on patterns of species diversity, we found that while the most parsimonious, ecologically coherent model of Australia’s biophysical dissimilarity (i.e., optimal partition of SAR)—like WWF’s biome map—also aggregated bioregions into seven groups, our algorithmically driven bioregionalization differed from the bespoke WWF’s expert-derived classification (Fig. 1 a). When assessing the ecological significance of the spatial configuration of these two biogeographic scenarios—based on a log-log implementation of the power function to fit the species-area relationship—we found that the increase in species richness with area varies among distinct spatial regions and taxonomic groups within regions (Fig. 1 b and c). Notably, similar—if not the same—patterns of species richness per standard area across taxonomic groups were detected between two regions of the optimal partition of SAR and those of WWF’s expert-derived classification that overlapped (i.e., Group 1 vs. Biome 4 - Temperate Broadleaf & Mixed Forests; and Group 3 vs. Biome 6 - Tropical & Subtropical Grasslands, Savannas & Shrublands), despite geographical differences due to the number of bioregions within their boundaries.

When hierarchical systems of IBRA bioregions were defined based on pairs of the best models of the variance in species richness and multiple-site compositional dissimilarity (i.e., optimal partitions of alpha and beta diversity), we found that like for the optimal partition of SAR, the distinct spatial clusters for the alpha- and beta-diversity of birds, mammals, herpetofauna, vertebrates and vascular plants largely included adjacent bioregions (Fig. S8.8 c– l), even with no explicit spatial aggregation or distance penalty being imposed on the algorithm. For vertebrate and vascular plant species, the spatial configuration and the numbers of optimal discrete clusters were the same for their alpha and beta diversity (Fig. S8.8 i–l)—with distinct plant-species areas almost perfectly collapsing within those for vertebrates (Fig. 2). Further, the distribution of the height of nodes in the species composition dendrograms was right skewed across all taxa, with the variation of compositional dissimilarity being larger for herpetofauna and vascular plants (Fig. 3).

**FIGURE 2.**
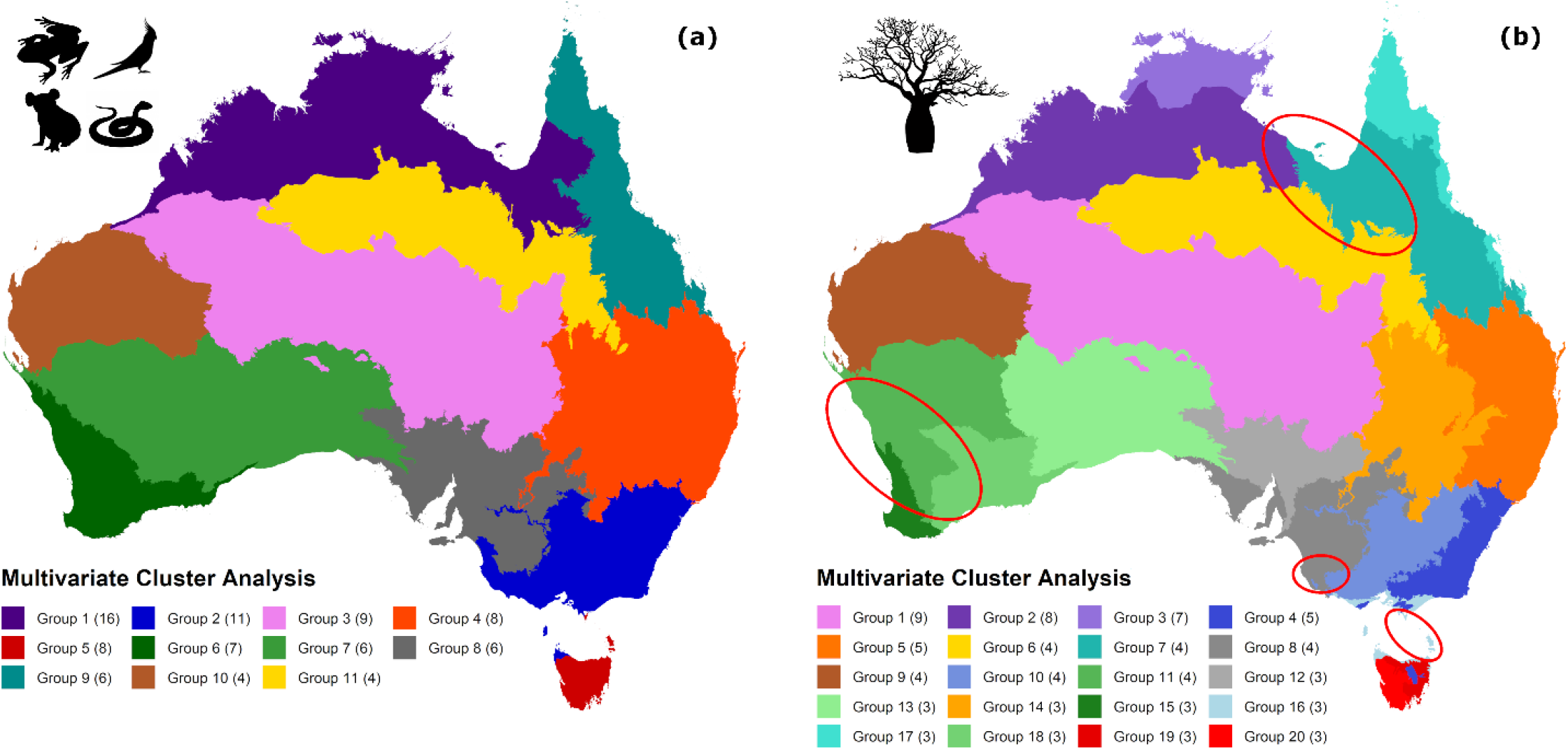
Spatial configuration of the optimal clusters for species diversity. (a) Vertebrate and (b) vascular plant species. Ellipses highlight the areas where plant clusters do not collapse within vertebrates. The number of bioregions in distinct spatial clusters are found between parentheses.

**FIGURE 3.**
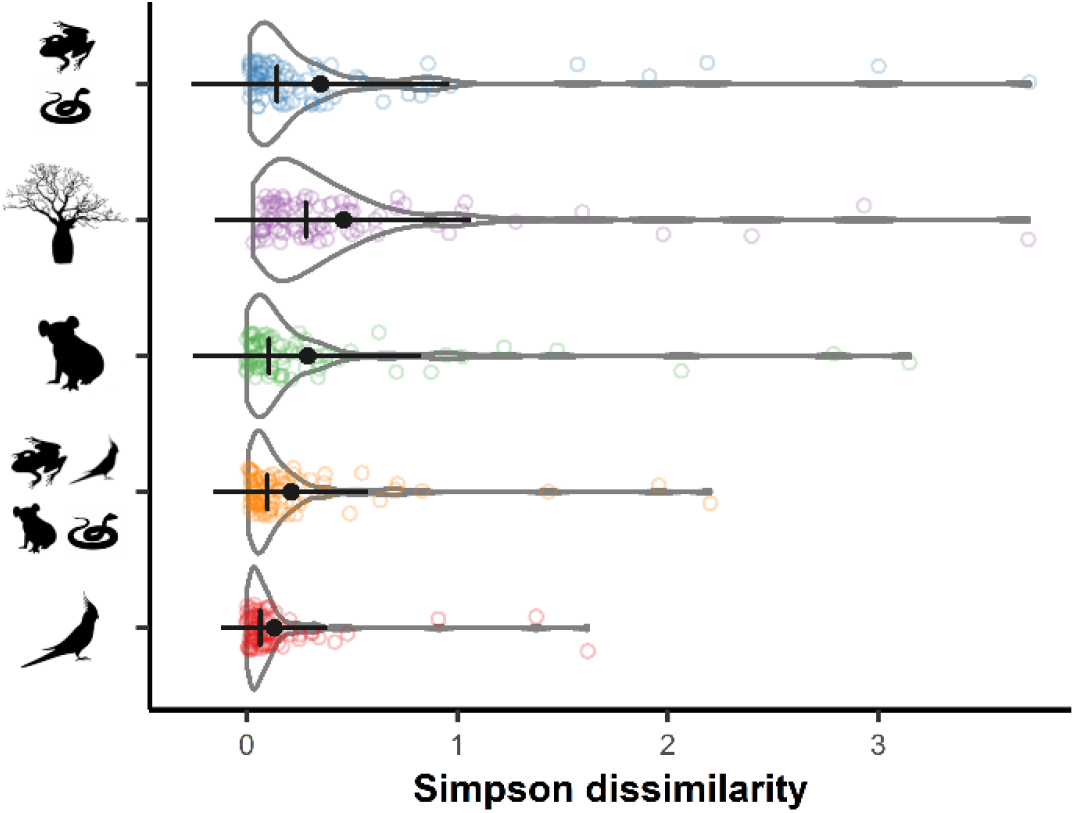
Height of nodes in five dendrograms. Kernel density plot shows the distribution of the nodes’ height for the spatial turnover of bird, mammal, herpetofauna, vertebrate, and vascular plant species. Color circle corresponds to the height of nodes as clusters are merged. Point and crossbar respectively represent the mean and median dissimilarity among nodes’ height, and dark line shows the standard deviation.

Within an information-theoretical analytical framework (Nowosad & Stepinski, 2018), the degree of spatial association between WWF’s biome map and those of optimal discrete-species clusters was moderate (Table 2), ranging from 50% of the information shared between the biome map and the spatial configuration of beta diversity in birds, and 58% for alpha and beta diversity in vascular plants. At the ecoregional level, WWF’s bioregionalization of Australia captured an additional 4 to 25% of information on patterns of species diversity embodied within our suite of optimal clusters, except when compared to the optimal discrete-bird map of beta diversity (i.e., − 4.3% spatial association between maps). The spatial concordance of optimal discrete-plant and -vertebrate clusters was slightly worse with each other than with other optimal discrete-species clusters at lower taxonomic ranks only for the pairwise-map comparisons between discrete-vertebrate and -mammal clusters (Table 2), for which the loss of information on alpha and beta diversity ranged from 0.8% to 4%, respectively. When contrasting the spatial association results of these two biogeographic scenarios, an algorithmically driven IBRA framework of discrete-plant and -vertebrate clusters retained more information on species alpha and beta diversity patterns within bioregions across multiple taxonomic groups than WWF’s hierarchical system of biomes and ecoregions. Nonetheless, the bespoke expert partitioning of Australia (IBRA) performed slightly better at retaining differences in species richness between bioregions of varying sizes (Table 2).

**Table 2.**
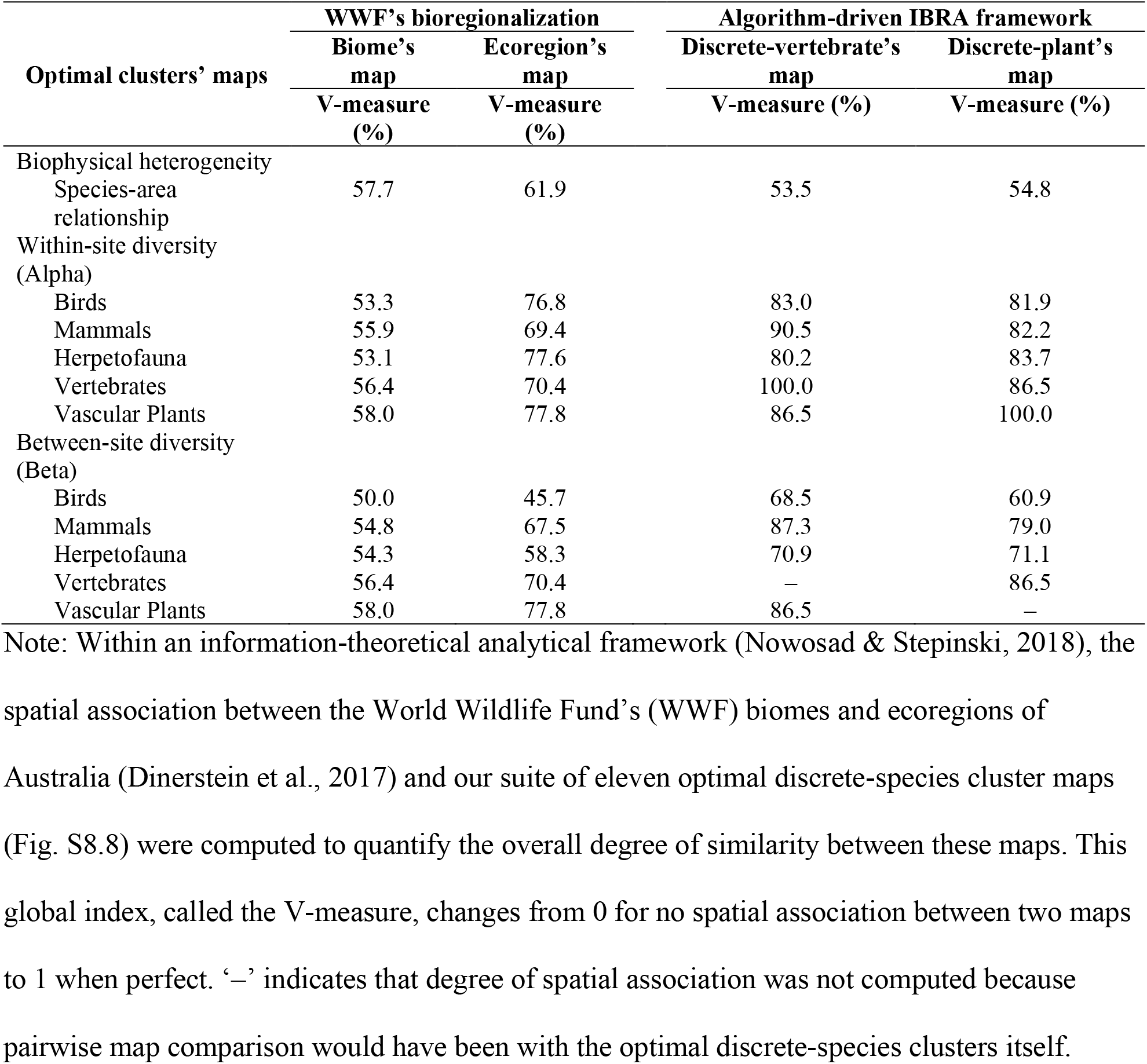
Mutual information on species richness and composition across pairwise-map comparisons.

| Optimal clusters' maps | WWF's bioregionalization |  | Algorithm-driven IBRA framework |  |
| --- | --- | --- | --- | --- |
|  | Biome's map | Ecoregion's map | Discrete-vertebrate's map | Discrete-plant's map |
|  | V-measure (%) | V-measure (%) | V-measure (%) | V-measure (%) |
| Biophysical heterogeneity |  |  |  |  |
| Species-area relationship | 57.7 | 61.9 | 53.5 | 54.8 |
| Within-site diversity (Alpha) |  |  |  |  |
| Birds | 53.3 | 76.8 | 83.0 | 81.9 |
| Mammals | 55.9 | 69.4 | 90.5 | 82.2 |
| Herpetofauna | 53.1 | 77.6 | 80.2 | 83.7 |
| Vertebrates | 56.4 | 70.4 | 100.0 | 86.5 |
| Vascular Plants | 58.0 | 77.8 | 86.5 | 100.0 |
| Between-site diversity (Beta) |  |  |  |  |
| Birds | 50.0 | 45.7 | 68.5 | 60.9 |
| Mammals | 54.8 | 67.5 | 87.3 | 79.0 |
| Herpetofauna | 54.3 | 58.3 | 70.9 | 71.1 |
| Vertebrates | 56.4 | 70.4 | — | 86.5 |
| Vascular Plants | 58.0 | 77.8 | 86.5 | — |

Note: Within an information-theoretical analytical framework (Nowosad & Stepinski, 2018), the spatial association between the World Wildlife Fund's (WWF) biomes and ecoregions of Australia (Dinerstein et al., 2017) and our suite of eleven optimal discrete-species cluster maps (Fig. S8.8) were computed to quantify the overall degree of similarity between these maps. This global index, called the V-measure, changes from 0 for no spatial association between two maps to 1 when perfect. '—' indicates that degree of spatial association was not computed because pairwise map comparison would have been with the optimal discrete-species clusters itself.

## 3. Discussion

Just as endemism is a commonly used basis for bioregionalizations (Ebach & Parenti, 2015; Morrone, 2014), species’ intrinsic traits are also responsible for defining ecologically meaningful clusters at large scale. We found strong spatial concordances across taxonomic groups and patterns of species richness and composition among the suite of optimal discrete-species clusters, signaling the effect that species’ biology and evolutionary history have in the identification of distinct areas of co-occurring species. The variation of compositional dissimilarity across our five target taxonomic groups (Fig. 3) reflects the overall small distributions of species with restricted range and/or low occupancy (Kreft & Jetz, 2010) and suggests that species’ dispersal abilities are important determinants of the emergent biogeographic divisions. Furthermore, having discrete plant-species areas almost perfectly nested within larger vertebrate-species areas suggests that interspecific relationships are also involved in explaining most of the variation in species diversity. Consequently, basing a hierarchical system of bioregions on plants and vertebrate optimal clusters is ecologically intuitive, because plants are essential to all animals, and interspecific interactions are largely responsible for generating community structure (Wisz et al., 2013).

Underpinning hierarchical systems of bioregions solely in the analysis of geographic and environmental covariates is not necessarily so strong as to capture the distributional patterns of multiple taxa in unison. Variation in the z-values (log-slopes) of the species-area relationship among discrete macrounits of biodiversity, such as the case of biomes, has already been documented (Kier et al., 2005). Yet, when combined with both the variability of z-values across taxonomic groups, and the parsimony of distinct spatial clusters for beta diversity with respect to those for alpha diversity (Fig. S8.8 c–l), it suggests that although environmental gradients are not impenetrable barriers to the arrangement of distinct communities, they can be important determinants of species richness within the structure of those communities. This demonstrates that while environmental heterogeneity is a well-established driver of species richness (Stein, Gerstner, & Kreft, 2014), defining a hierarchical system of bioregions that meaningfully aggregates biodiversity should not rely on species richness alone, because it varies among higher taxonomic groups, and does not necessarily have a positive relationship with species endemism (Koleff & Gaston, 2002).

Considering that WWF’s biomes were defined using associations of climate and dominant vegetation forms and structure to broadly classify terrestrial ecosystems (Kier et al., 2005; Mackey et al., 2008), the relatively moderate degree of the overall spatial association between WWF’s biomes and our eleven optimal-cluster maps was expected. This finding aligns qualitatively with those reported in recent studies (Coops et al., 2018; Edler et al., 2017), and they together give strength to our conclusion that complex interactions between biophysical factors and species’ intrinsic attributes are reflected in a nested hierarchy of bioregions (Sexton, McIntyre, Angert, & Rice, 2009). This means that hierarchical bioregionalization systems must account for variability in the distributional patterns of different taxa (Morrone, 2018) if they are to direct more efficient and appropriately targeted biodiversity research, policy, and conservation across multiple spatial or temporal planning scales.

Bias in biodiversity conservation is systemic (Butchart et al., 2015) and usually attributed to conservation efforts being implemented, for practical reasons, in areas of low agricultural productivity (Joppa & Pfaff, 2009) and/or targeting specific groups of species, such as charismatic species (Colléony, Clayton, Couvet, Saint Jalme, & Prévot, 2017). However, biogeographic frameworks might also contribute to this bias. When comparing WWF’s hierarchical system (ecoregions nested within biomes) to an algorithmic, data-driven hierarchical system of IBRA bioregions based on plant- and vertebrate-oriented optimal areas, the difference in the amount of information on Australian biodiversity patterns that each tier of these two bioregionalization scenarios captured was greater for the (bespoke) WWF’s expert-derived classification, with neither biomes nor ecoregions outperforming an integrative, data-driven alternative at retaining information on alpha and beta diversity across taxa (Table 2). This finding suggests that—from inception—systematic conservation planning may have been undermined by how a particular biogeographic framework was defined, which in the case of WWF’s global bioregionalization included biotopic and biocoenotic classification approaches (Mackey et al., 2008). This is troublesome if we consider that, in addition to being one of the most widely used biogeographic templates for biodiversity conservation, WWF’s ecoregions have served as the basis of other well-established conservation strategies at global scale (Lamoreux et al., 2006). Given that spatial information on biodiversity patterns is essential for effective biodiversity conservation, the design of environmental policies, the establishment of protected area networks, and the implementation of more recent *in situ* interventions (e.g., rewilding, species’ translocations), there is a substantial need for a hierarchical system of geographic operational units that is ecologically interpretable across broad spatial and taxonomic breaths.

Over the past 200 years, the history of bioregionalization in Australia has been driven as much by changes in foci (i.e., from exploration, to the conservation of biodiversity) as by theoretical and methodological advances and data availability (Ebach, 2012). Multiple studies have found that different biogeographic templates, encompassing those both qualitative and quantitative perspectives, were mostly congruent with each other (Bloomfield, Knerr, & Encinas-Viso, 2018; González-Orozco, Laffan, Knerr, & Miller, 2013; González-Orozco, Thornhill, Knerr, Laffan, & Miller, 2014). Yet, even when applying similar biogeographic approaches to partition Australia’s landscape, phyto- and zoo-geographers disagreed on the boundaries of some areas of Australia, such as the arid region (Ebach & Murphy, 2020). In our algorithmic, data-driven model, the partition of arid Australia is greater than in any of the bioregionalization scenarios of Bloomfield et al. (2018). Yet, despite methodological differences, a visual comparison of these two approaches’ bioregionalization scenarios suggests that geographic operational units of optimal discrete-vertebrate and -plant clusters in arid Australia are spatially nested within the zones delineated in the study of Bloomfield et al. (2018), which the authors in turn argued to be spatially consistent with Eremaean biogeographic region (i.e., arid Australia based on its flora).

As a basis for developing ecologically sensible bioregionalizations that are both methodologically robust and repeatable, we argue that this new approach is highly innovative and can be applied in many contexts. Nonetheless, there are some caveats. First, no consensus exists on how to decisively identify biogeographic divisions and to delineate their boundaries (Antonelli, 2017; Morrone, 2018). Yet, the plethora of aggregative approaches to emerge in biogeographic research has been instrumental in our understanding that any single bioregionalization cannot hope to consistently capture the distributional patterns of multiple taxa (Coops et al., 2018; Edler et al., 2017; Kreft & Jetz, 2010; Vilhena & Antonelli, 2015). While our study reinforces this conclusion, we do show that a hierarchical system more directly relevant to biodiversity can be derived systematically by leveraging information on species diversity and composition within a bioregionalization’s geographic operational units. Second, biased, and inadequate knowledge on species’ distribution can confound the identification of natural biogeographic areas (Pimm et al., 2014; Whittaker et al., 2005). However, a hierarchical bioregionalization system based on our algorithmic approach can be readily and transparently revised to reflect the increasing availability of distributional data, and the species’ responses to global and regional environmental changes. Third, because diversity is unevenly distributed between taxa, as well as in space and time (Whittaker et al., 2005) other biodiversity dimensions beyond species might be relevant. As functional and phylogenetic data continue to accumulate, and comparative approaches are advanced, a similar approach might be used to investigate whether a general bioregionalization is maintained under such patterns, and how the different dimensions of biodiversity interrelate within a particular biogeographic framework.

This work demonstrates how a systematic, objective examination of the patterns of species diversity within the geographic operational units of a biogeographic template—in this study, 85 IBRA bioregions—can be used to develop a rigorous hierarchical system of discrete spatial partitions that is directly relevant for aggregating biodiversity. The use of existing, geographically restricted operational units, which might also be delineated using quantitative techniques, makes our approach not only generic (Mackey et al., 2008), but also sufficiently flexible to account for the increasing knowledge of biodiversity. As such, this robust, data-driven method can underpin the design and implementation of *in situ* conservation initiatives in other regions and even globally, inform policy for meaningful environmental and biodiversity outcomes across multiple planning scales, and bolster interests in the factors and the processes driving discontinuities of biodiversity. This is crucial, given the effect of human-mediated habitat transformations on the boundaries of biogeographical regions globally.

## Data availability statement

The data on climate, elevation, vegetation, soil, lithology, and the subregions of the Interim Biogeographic Regionalization for Australia (IBRA) framework are freely available online. All the sources are cited in the manuscript. The custom computer code used for this study, as well as the processed spatial information for subregions and bioregions of the IBRA framework, the species occurrences post-equalization, the geographic-based measures of biophysical factors, the presence-absence matrices for vascular plants and four vertebrate classes (amphibians, birds, mammals, and reptiles), and the information on all fitted models for the relationship between species richness and size of IBRA geographic operational units are freely available at: [DOI will be provided should the manuscript be accepted for publication].

## Appendices

Appendix S1. Additional information on data collection and processing

Appendix S2. Selection of species-area relationship (SAR) mathematical function

Appendix S3. Selection of biophysical factors and IBRA unit of analysis

Appendix S4. Selection of optimal discrete-species clusters

Figs. S5 to S8

List of references

