## Supplemental Information for "Robust, data-driven bioregionalizations emerge from diversity concordance"

**This document includes:**

Appendix S1. Additional information on data collection and processing

Appendix S2. Selection of species-area relationship (SAR) mathematical function

Appendix S3. Selection of biophysical factors and IBRA unit of analysis

Appendix S4. Selection of optimal discrete-species clusters

Figs. S5 to S8

List of references

**Appendix S1. Additional information on data collection and processing**

**Spatial data**

In version 7 of the IBRA framework (Department of Agriculture, 2012b), 89 bioregions are further divided in 419 subregions. From the shapefile representing IBRA, we omitted all polygons that correspond to oceanic islands and islets (< $1km^2$ landmass). This resulted in the removal of four IBRA bioregions (i.e., Coral Sea, Indian Tropical Islands, Pacific Subtropical Islands, and Subantarctic Islands) and one subregion (i.e., Southern Great Barrier Reef) nested within South Eastern Queensland IBRA bioregion.

We used this spatial information (n = 1,362 discrete polygons) to derive a four-tier hierarchical system of geographic operational units for Australia—where 410 IBRA subregions are nested within 85 IBRA bioregions, which are nested within 37 ecoregions, and those macrounits are themselves embedded in 7 broader-scale (and spatially coherent) biomes based on the most recent version of WWF's bioregionalization (Dinerstein et al., 2017). This was possible as bioregions are a more detailed geographic classification of the Word Wildlife Found (WWF) Terrestrial Ecoregions (Department of Agriculture, 2012a). Our goal in doing this was to base our analyses on the most up-to-date bioregionalization for the conservation of Australia's biodiversity, from regional-to-global scale, and to account explicitly for the possible confounding effect that species endemism in small islands (Lomolino & Weiser, 2001; Triantis & Sfenthourakis, 2012) and mapping errors between tiers might exert on the biophysical and biotic characterization of mainland Australia and Tasmania.

**Climatic variables**

We used all nineteen climatic variables generated for Australia by the Biodiversity and Climate Change Virtual Laboratory (Hallgren et al., 2016). This includes Annual mean temperature, Mean diurnal range, Isothermality, Temperature seasonality, Maximum temperature of warmest month, Minimum temperature of coldest month, Temperature annual range, Mean temperature of wettest quarter, Mean temperature of driest quarter, Mean temperature of warmest quarter, Mean temperature of coldest quarter, Annual precipitation, Precipitation of wettest month, Precipitation of driest month, Precipitation seasonality, Precipitation of wettest quarter, Precipitation of driest quarter, Precipitation of warmest quarter, and Precipitation of coldest quarter.

### Excluded categories from discrete variables

We excluded 7 of 33 major vegetation groups because of their extremely restricted geographic extent (i.e., Unclassified forest; and Unclassified native vegetation) and/or inconsequence for characterizing terrestrial native vegetation (i.e., Cleared non-native vegetation buildings; Inland aquatic – freshwater, salt lakes, lagoons; Naturally bare sand rock claypan mudflat; Sea and estuaries; and Unknown/no data). Based on these arguments, we also excluded one out of thirteen lithology classes (Unknown), and two out of fifteen soil types (No data; Rock).

### Quality control of Atlas of Living Australia records

Biases and errors are common issues in virtually all distributional datasets, which have been (and will continue to be) the basis of contemporary grid-based bioregionalizations. As in a previous study on the IBRA framework (Ondei, Brook, & Buettel, 2019), we excluded records that lacked geographic information, as well as those that were from before 1990 or classified as weeds (Randall, 2007) or exotic vertebrates (Vertebrate Pests Committee, 2007) in Australia. We omitted species occurrences with unidentifiable scientific names or those situated outside the geographic extent of the model system, resulting in a final dataset of 25,995 native species: 23,248 vascular plants, 233 amphibians, 1,201 birds, 349 mammals, and 964 reptiles.

### Assembly of species' distributional data

Kernel functions are a well-established technique to estimate the area of occupancy of a species, but this method relies on a homogeneous sampling effort across the landscape to produce reliable results. To meet this requirement, we developed a procedure to even out the difference in sampling effort (i.e., number of species occurrences) among subregions by minimizing the variance within subregions' size-classes, while maximizing the variance between size-classes. In this equalization procedure, we used, as a base of comparison, the mid-point (mean) of the Jenks natural-breaks optimization (Jenks, 1977; Jenks & Caspall, 1971) of IBRA subregions' log area ($km^2$) into three size-classes (small, medium, and large) to thin the number of occurrences within subregions based on a random selection.

For species with 20 or more occurrences post-equalization, we derived extent-occurrence maps that captured 99% of species' empirical distribution in mainland Australia and Tasmania using a Gaussian (bivariate normal) kernel density function with plug-in estimator as smoothing parameter. To avoid overestimating a species' area of occupancy, we chose to do this separately for mainland Australia and Tasmania based on whether species records were from either of these two landmasses. Additionally, we cropped the empirical extent-occurrence maps for mainland Australian and Tasmanian species to the bounding-box of their corresponding geographic extent to avoid the blending of endemic species between these two landmasses. With this spatial stratification for deriving extent-occurrence maps, we sought to account for the phenomenon that patterns of species diversity are unevenly distributed in space and time, as well as across taxa (Jetz, McPherson, & Guralnick, 2012; Whittaker et al., 2005).

In addition to homogenizing sample effort across IBRA operational units (i.e., equalization procedure), selecting the smoothing parameter is crucial for the estimation of utilization distributions (Epanechnikov, 1969). We chose this habitat-utility framework as the best-suited alternative (i.e., less prone to overestimating species empirical distributions) for this dataset after visually comparing the extent-occurrence maps for a set of cosmopolitan, coastal, and rare bird species with 95% and 99% confidence region to these species' extent-occurrence maps, derived using reference bandwidth as smoothing parameter instead (Fig. S1.1). The selected framework proved to be suitable for a set of extent-occurrence maps for amphibians, mammals, reptiles, and plants endemic to mainland Australia and/or rare to Tasmania (Fig. S1.2). We thus applied it to all investigated taxa.

In this study, we considered a species to be rare/range-restricted when it had less than 20 occurrences post-equalization, matching the theoretical minimum sample size needed to ensure that the relative mean square error is not greater than 0.1 when estimating a standard bivariate density using a kernel function (Silverman, 1986). We used species occurrences with less than 20 records post-equalization as source of distributional information on species with potential narrow geographic range and/or low occupancy. Although this may result in the omission of certain species from some IBRA geographic operational units, their large size (e.g., the smallest IBRA subregion is ~ 200 $km^2$) minimizes the impact of this error (false absences) on our analyses, while including the whole pool of Australian native species.

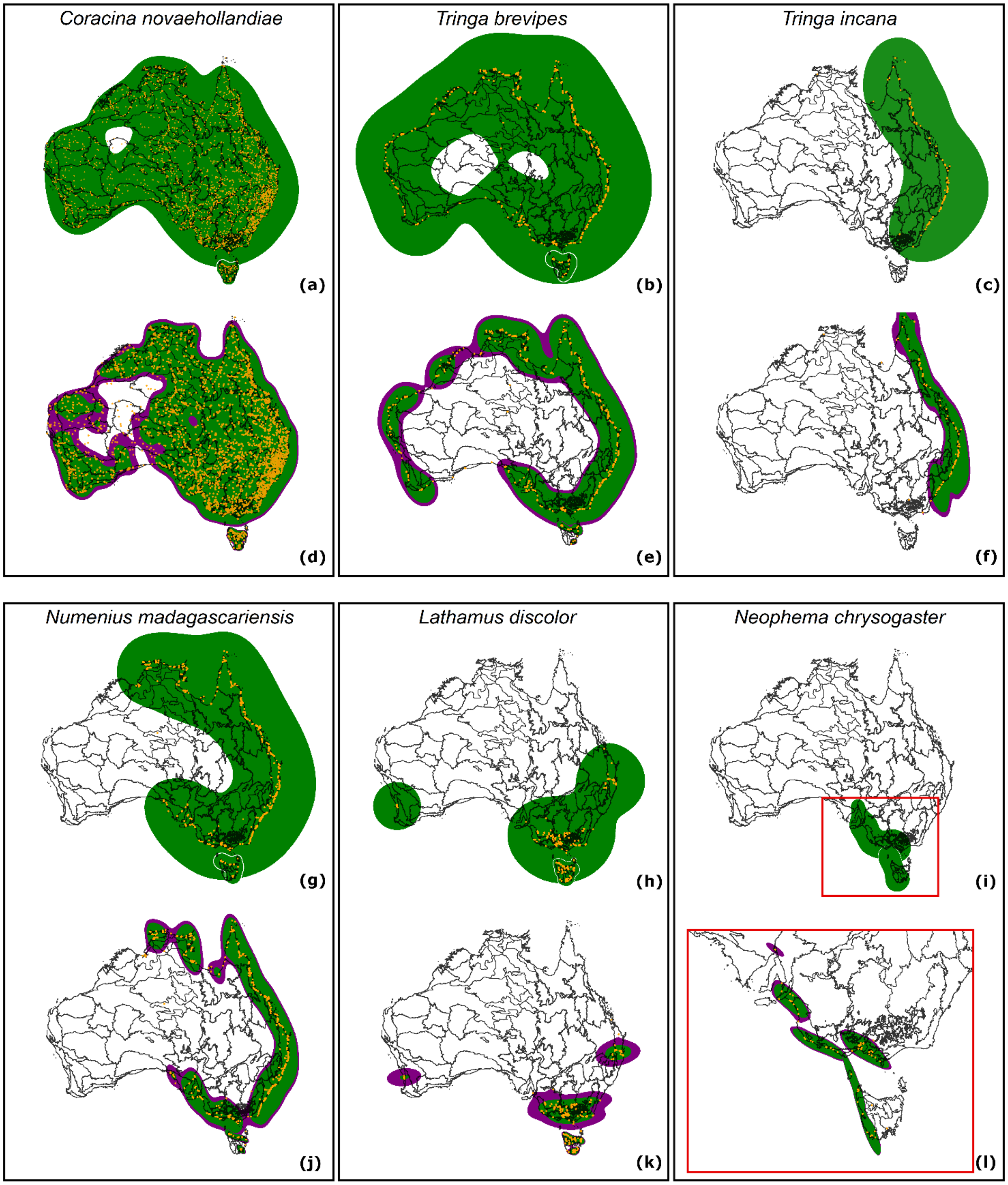


**FIGURE S1.1 Extent-occurrence maps for a suite of cosmopolitan, coastal, and rare bird species.** Overlap between Australia's bioregions and species' utilization distributions using a Gaussian (bivariate normal) kernel density function with (a–c, and g–i) the reference bandwidth or (d–f, and j–l) the plug-in estimator as smoothing parameter. Orange points corresponds to species occurrences post-equalization. Green and purple polygons respectively represent 95% and 99% confidence region of species' utilization distributions in Mainland and/or Tasmania. Polygons' grey outline (a, b, g, and h) delimits species' utilization distributions for Tasmania. Oceanic cells that fell within the confidence regions where subsequently excluded.

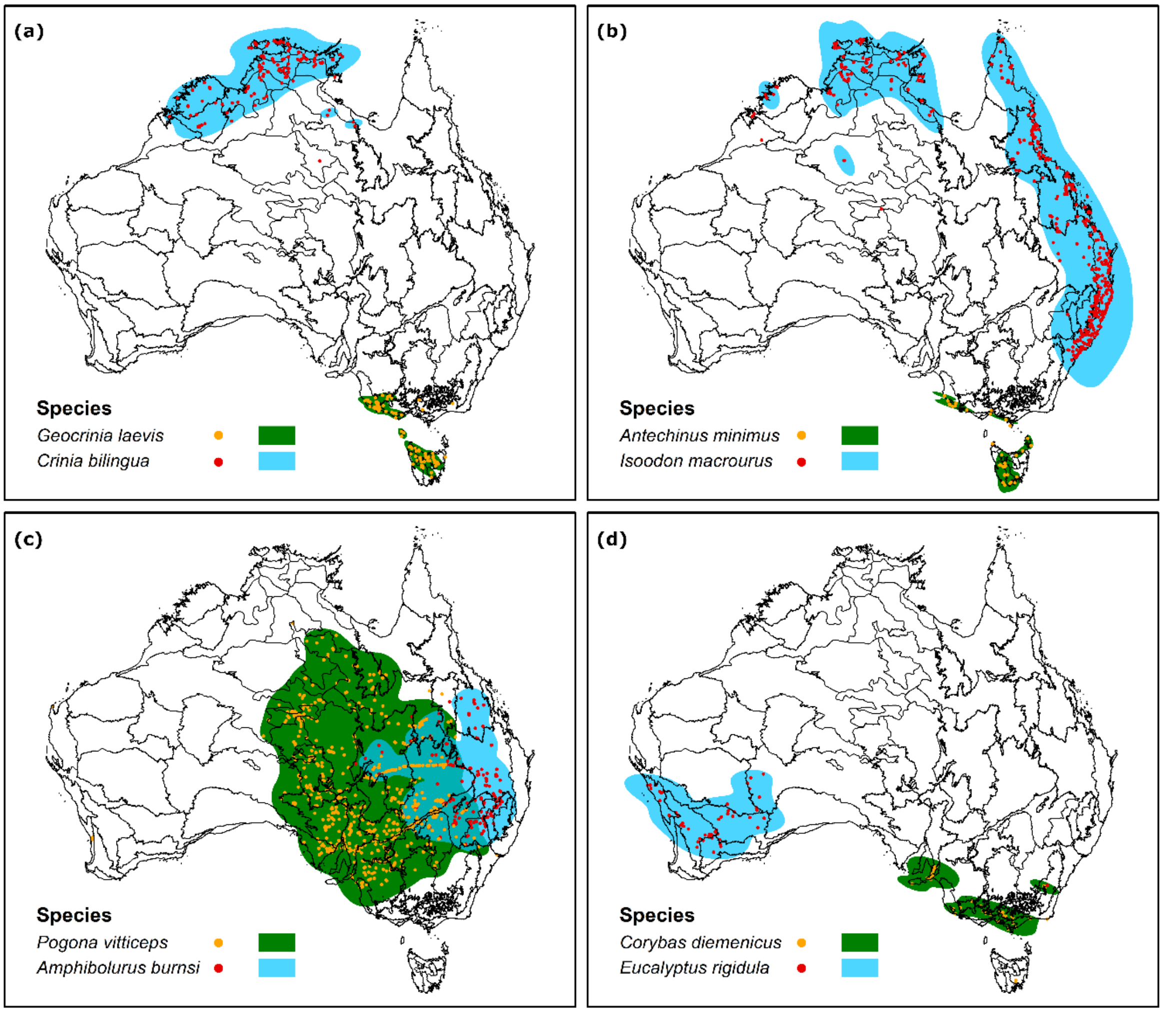


**FIGURE S1.2 Extent-occurrence maps for a suite of plant and selected vertebrate species.** Overlap between Australia's bioregions and species' utilization distributions using a Gaussian (bivariate normal) kernel density function with the plug-in estimator as smoothing parameter. Color points correspond to species occurrences post-equalization for (a) amphibians, (b) mammals, (c) reptiles, and (d) vascular plants. Color polygons correspond to 99% confidence region of species' utilization distributions. Oceanic cells that fell within the confidence regions where subsequently excluded.

Given the impracticality of estimating habitat suitability for all species (n = 25,995), combining species occurrences and empirical extent-occurrence maps represented an appropriate next-best option to base the estimation of within- and between-site diversity of species (alpha and beta). Like expert-based maps (Ficetola et al., 2014; Jetz, Sekercioglu, & Watson, 2008), which are commonly used in biogeographic and macroecological studies, our set of empirical extent-occurrence maps may incur commission errors (false presences) (Brooks et al., 2019). Yet, because these maps were based on the probabilistic distribution of the optimum number of species with respect to the size of subregions, they vary little in quality and may better reflect recent range contractions due to environmental change—two common issues of expert-based distributional maps (Di Marco, Watson, Possingham, & Venter, 2017; Rondinini et al., 2011)—as well as being reproducible and appropriate for dealing with outliers, such as vagrant individuals, and misidentifications.

### GIS and R packages

We used ArcGIS v. 10.5.1 (2017) to remove polygons of oceanic and small islands. Spatial calculations, and feature engineering were done in the program R v. 3.6.3 (R Core Team, 2020) using functions from the 'raster' v. 2.8-19 (Hijmans, 2019), 'spatialEco' v. 1.2-1 (Evans, 2019), 'sf' v. 0.1-7 (Pebesma, 2018), 'parallel' v. 3.6.3 (R Core Team, 2020), 'future' v. 1.14.0 (Bengtsson, 2019), 'tidyverse' v. 1.3.0 (Wickham et al., 2019), 'reshape2' v. 1.4.3 (Wickham, 2007), 'lwgeom' v. 0.1-7 (Pebesma, 2019), and 'magrittr' v. 1.5 (Bache & Wickham, 2014) packages.

In addition to the R packages listed above, we used 'BAMMtools' v. 2.1.6 (Rabosky et al., 2014) package to computed Jenks natural breaks of subregions' sizes, and 'ks' v. 1.11.5 (Duong, 2019), 'rgdal' v. 1.4-3 (Bivand, Keitt, & Rowlingson, 2019), 'maptools' v. 0.9-8 (Bivand & Lewin-Koh, 2019), 'gpclib' v. 1.5-5 (Peng, Murdoch, Rowlingson, & Murta, 2013), and 'PBSmapping' v. 2.72.1 (Schnute, Boers, & Haigh, 2019) packages to estimate species' empirical distributions and to convert the results to polygon features. Meanwhile, we used the 'adehabitatHR' v. 0.4.18 (Calenge, 2006) package to estimate the utility distributions for the alternative extent-occurrence maps of cosmopolitan, coastal, and rare bird species.

**Appendix S2. Selection of species-area relationship (SAR) mathematical function**

While the logarithmic form of the power function has been most frequently applied for fitting SAR (Tjørve, 2003; Triantis, Guilhaumon, & Whittaker, 2012), it is still debatable what mathematical function best describes this relationship (Scheiner, 2003). Hence, we fitted the log-log implementation of the power function alongside two other functions where the relationship was assumed linear (Power in log-log space through origin, and simple linear model in the arithmetic space), three functions where the shape of the curve was assumed convex (Exponential, Power, and Negative Exponential), and three other functions where the relationship was assumed sigmoid (Cumulative Weibull Distribution, Logistic, and Morgan-Mercer-Flodin). The last six mathematical functions are as in Triantis et al. (Triantis et al., 2012).

As a base of comparison, we used the expert-based classification of IBRA subregions into biomes to assess/rank these nine functions to model SAR for five taxonomic groups (bird, mammal, herpetofauna, vertebrate and vascular plants). The species richness and size of subregions were computed from presence-absence matrices, and subregions' polygon geometry, respectively. We omitted SAR models for the Montane Grasslands and Shrublands biome because its classification included only two subregions. The 'stats' v. 3.6.3 (R Core Team, 2020), and the 'sars' v. 1.2.0 (Matthews, Triantis, Whittaker, & Guilhaumon, 2019) packages were used to fit SARs when the relationship was assumed linear, and convex or sigmoid, respectively. To select the best function to model SAR, we used Akaike's Information Criteria (Akaike, 1974).

We found that SAR models based on the canonical log-log implementation of the power function outperformed all other models where the mathematic function used to fit the increase in the number of species per standard area was assumed convex or sigmoid. Notably, the SAR models based on the power in log-log space through origin were the second-best model for all cases. Forcing the intercept to pass through zero is not recommended. This result suggests that there exists a negative correlation between species-area ratio and area size (i.e., non-linearity in SAR; Ovadia, 2003) that cannot be detected with the richness-area structure of our datasets (i.e., number of species within IBRA subregions and their size in $km^2$). While not without its own limitations (Guilhaumon, Gimenez, Gaston, & Mouillot, 2008), we deemed the log-log implementation of the power function to be most appropriate model of SAR to examine patterns of species richness with increasing size of IBRA biogeographic conservation units.

### Appendix S3. Selection of biophysical factors and IBRA unit of analysis

Multiple Factor Analysis (MFA) revealed that 90% of the variance in the biophysical dataset at subregional and bioregional scale can be explained with 37 and 27 principal components, respectively. We found that the correlation between all five groups of variables and the first two dimensions were qualitatively similar in their contribution to describing the biophysical space of subregions and bioregions (Fig. S3.3 a and b). Only mean elevation contributed the least when changing to a coarser landscape resolution.

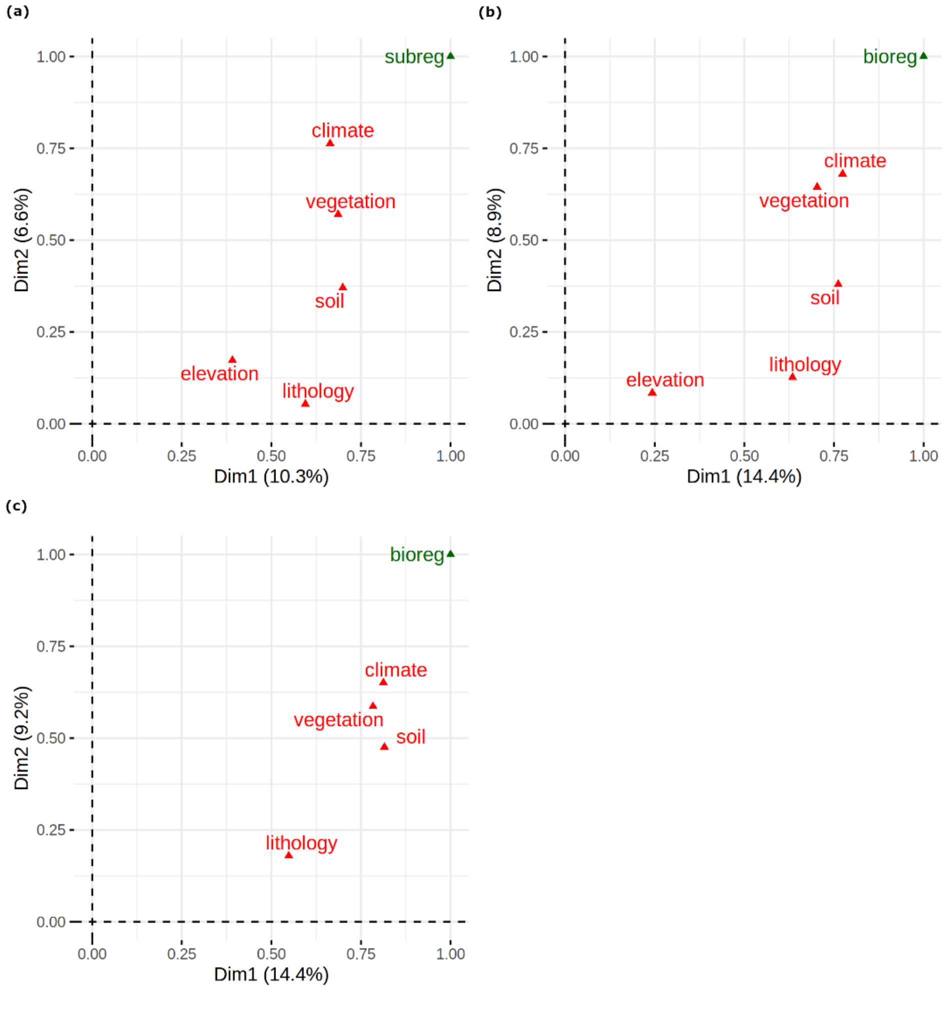


**FIGURE S3.3 Multiple factor analysis (MFA) of Australia's biophysical space.** The first two principal components of the global multidimensional biophysical space for the geographic operational units of the Interim Biogeographic Regionalization for Australia (IBRA) framework. (a–b) MFA of a set of five geographic and environmental covariates (i.e., climate, elevation, lithology, soil, and vegetation) computed for subregions (subreg) and bioregions (bioreg) of the IBRA framework, respectively. (c) MFA of a set of four geographic and environmental covariates (climate, lithology, soil, and vegetation) computed for bioregions.

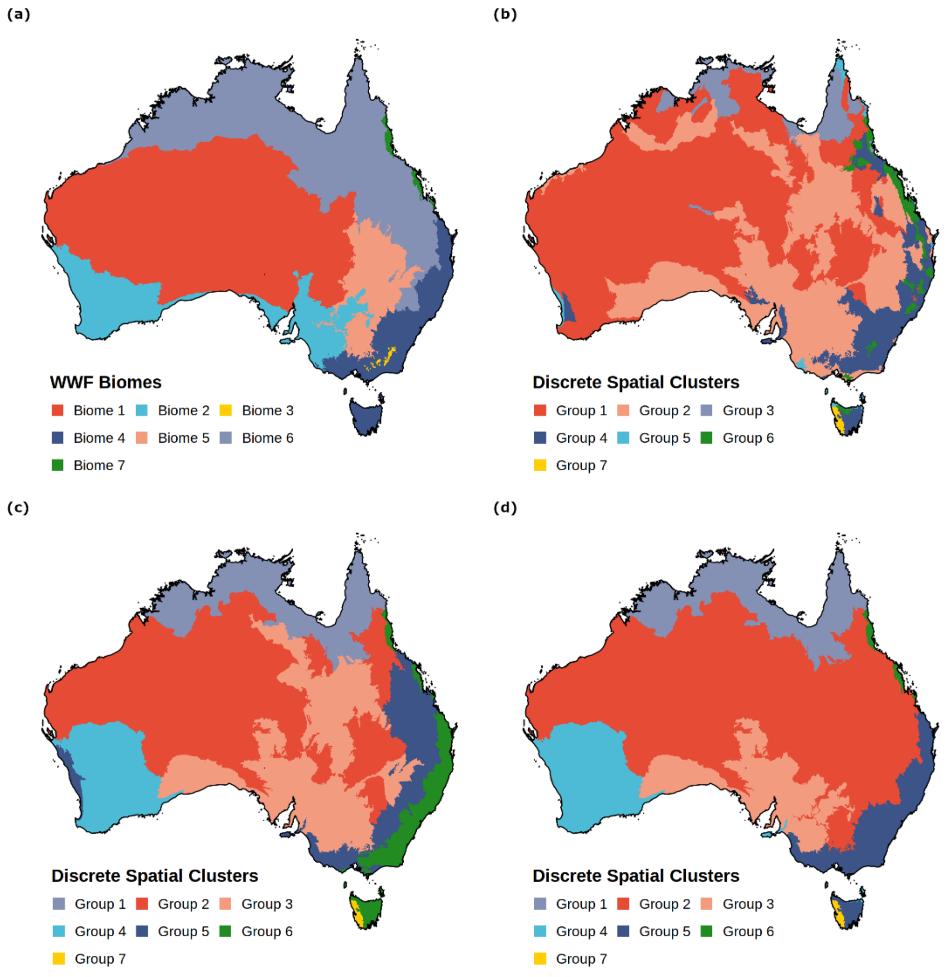


**FIGURE S3.4. Spatial nature of dissimilarity of Australia's biophysical space.** Maps corresponding to (a) the World Wildlife Fund's (WWF) expert classification of Australia into seven biomes (Dinerstein et al., 2017) or the algorithm-driven aggregation of (b) subregions and (c–d) bioregions of the Interim Biogeographic Regionalization for Australia framework into seven groups using the principal components of multiple factor analysis of either (b and c) five (climate, elevation, lithology, soil, and vegetation); or (d) four (climate, lithology, soil, and vegetation) suites of geographic and environmental covariates. Biome 1 = Deserts & Xeric Shrublands, Biome 2 = Mediterranean Forests, Woodlands, Scrub, Biome 3 = Montane Grasslands & Shrublands, Biome 4 = Temperate Broadleaf & Mixed Forests, Biome 5 = Temperate Grasslands, Savannas & Shrublands, Biome 6 = Tropical & Subtropical Grasslands, Savannas & Shrublands, and Biome 7 = Tropical & Subtropical Moist Broadleaf Forests. Color codes reflect the general geographic distribution of groups of bioregions in the Australian landscape, although—where possible—colors match those used to differentiate WWF's biomes.

The spatial assessment of these two global analyses differed greatly, with an increase in spatial coherence and ecological interpretability of the seven clusters when the ordination was based on principal components at bioregional level (Fig. S3.4 b and c). These findings signal bioregions as the most appropriate level for biodiversity analyses within the IBRA framework, although the remains noise in the structure of the biophysical dataset. Consequently, we performed an additional MFA of bioregional Australia without elevation for a subsequent spatial assessment (Fig. S3.3 c). Based on 27 principal components, we found that bioregions defined algorithmically in this way were coherently aggregated, and that their ordination aligned closer to the bespoke expert-based knowledge approach (Fig. S3.4 a and d). This confirms bioregions as the appropriate unit of analysis and reduces the biophysical dataset’s structure to include the eigenvalues of the 27 principal components based on climate, lithology, soil, and vegetation information in all subsequent analyses.

### Appendix S4. Selection of optimal discrete-species clusters

#### Process to identify and discriminate suites of prominent clusters

For the dendrogram in the biophysical space, we used the 'L' technique (Salvador & Chan, 2004) to identify the point of maximum curvature based on the evaluation plot of the merging height of nodes in the dendrogram versus the number of clusters. This intersection point represented the model that most accurately fitted the dataset while also minimizing complexity. We used this result to define the ceiling in the numbers of groups that prominent clusters using the 'dynamic' technique (Langfelder, Zhang, & Horvath, 2008) should not exceed, with the outputs of the 'dynamic' technique subsequently also defining the desired number of clusters for the 'static' technique.

We selected the optimal partition by examining both the spatial coherence and ecological significance of this set of prominent clusters, alongside the expert-based classification of bioregions into 7 biomes based on a log-log species-area relationship for our five target taxonomic groups. When the expert-based classification of bioregions into biomes or prominent clusters included groups with less than three bioregions—such as was the case for the Montane Grasslands and Shrublands, and the Tropical and Subtropical Moist Broadleaf Forests biomes—they were not considered in the identification of best SAR (i.e., species-area relationship) models. While neither slope nor standard error of SAR models could be computed for those restricted-group cases, the slope was estimated when the bioregionalization included groups with at least two bioregions, such as was the case for the Tropical and Subtropical Moist Broadleaf Forests biome. We used R-squared to measure the amount of variance explained by SAR models, and AIC (i.e., Akaike's Information Criteria) to select the most parsimonious SAR models.

Using both the 'dynamic' and 'static' techniques, we cut the dendrograms of compositional dissimilarity to define the set of prominent clusters for birds, herpetofauna, mammals, vertebrates, and vascular plants. In contrast to the stepwise process used to define prominent clusters of biophysical factors, these sets of prominent clusters included all five results of the 'dynamic' technique, and the results when using the 'static' technique to cut dendrograms into 2 to 37 groups. For all cases, we estimated the within- and across-variance of alpha and beta diversity by fitting ANOVA models of species richness, and by the sum of multiple-site measures of compositional dissimilarity across groups of bioregions in the set of prominent clusters divided by the multiple-site measure of compositional dissimilarity across Australia's bioregions, respectively. We assessed the relevance of these sets of prominent clusters by inspecting evaluation plots.

For alpha diversity, we plotted AIC and the Bayesian Information Criteria (BIC) (Schwarz, 1978) scores against the number of clusters. The optimal partitions for alpha diversity were selected from the tree-cutting technique with the clearest break in the maximum likelihood estimation of the regression line based on whether AIC and BIC scores agreed on the numbers of clusters. When agreement between both penalized-likelihood criteria was not achieved, the prominent clusters with the lowest BIC score was selected as optimal. This is because in terms of sensitivity and specificity BIC is more appropriate in situations where a false positive finding (Type I error) is considered as or more, misleading than its false negative (Type II error) counterpart (Dziak, Coffman, Lanza, Li, & Jermiin, 2012, 2020).

Meanwhile, we plotted the ratio of multiple-site measures of compositional dissimilarity in each of our suite of prominent clusters and that of Australia against the number of clusters to assess the relevance of prominent clusters for beta diversity. In this case, the prominent cluster with the highest ratio–which corresponds to the highest amount of variation in species composition–was selected as optimal for each taxonomic group. When the goal is to compare the variation in species composition across a set of sites, multiple-site measures of dissimilarity are strongly recommended due to the incapacity of averaged pairwise dissimilarity indexes of reflecting the multiple-site nature of dissimilarity (Baselga, 2012). Yet, these measures suffer from a site-number dependency (Baselga, 2010). To account for this, we calculated the average of multiple-site measures based on a resampling procedure iterated 100 times, and in which the minimum number of bioregions among the groups in each prominent cluster defined its sample size.

Multiple-site measures of compositional dissimilarity were computed using the 'betapart' v. 1.5.1 (Baselga & Orme, 2012) package. To cut dendrograms based on the 'dynamic' technique, we used cutreeDynamic function in the 'dynamicTreeCut' v. 1.63-1 (Langfelder et al., 2008) package, with the method set to hybrid, the minimum number of cluster size set to one, and the relative sensitivity to cluster splitting set to five default parameters. We used the 'stats' v. 3.6.3 (R Core Team, 2020) package to estimate the within- and across-variance of alpha diversity, and to define clusters based on the ‘static’ tree-cutting technique. We used the ‘broom’ v. 0.5.4 (Robinson & Hayes, 2020) package to extract summaries on the goodness of fit (R-squared, AIC, and/or BIC values) of the within- and across-variance in alpha diversity, and of the species-area relationship, as well as on the components of fitted SARs (slope, and standard error).

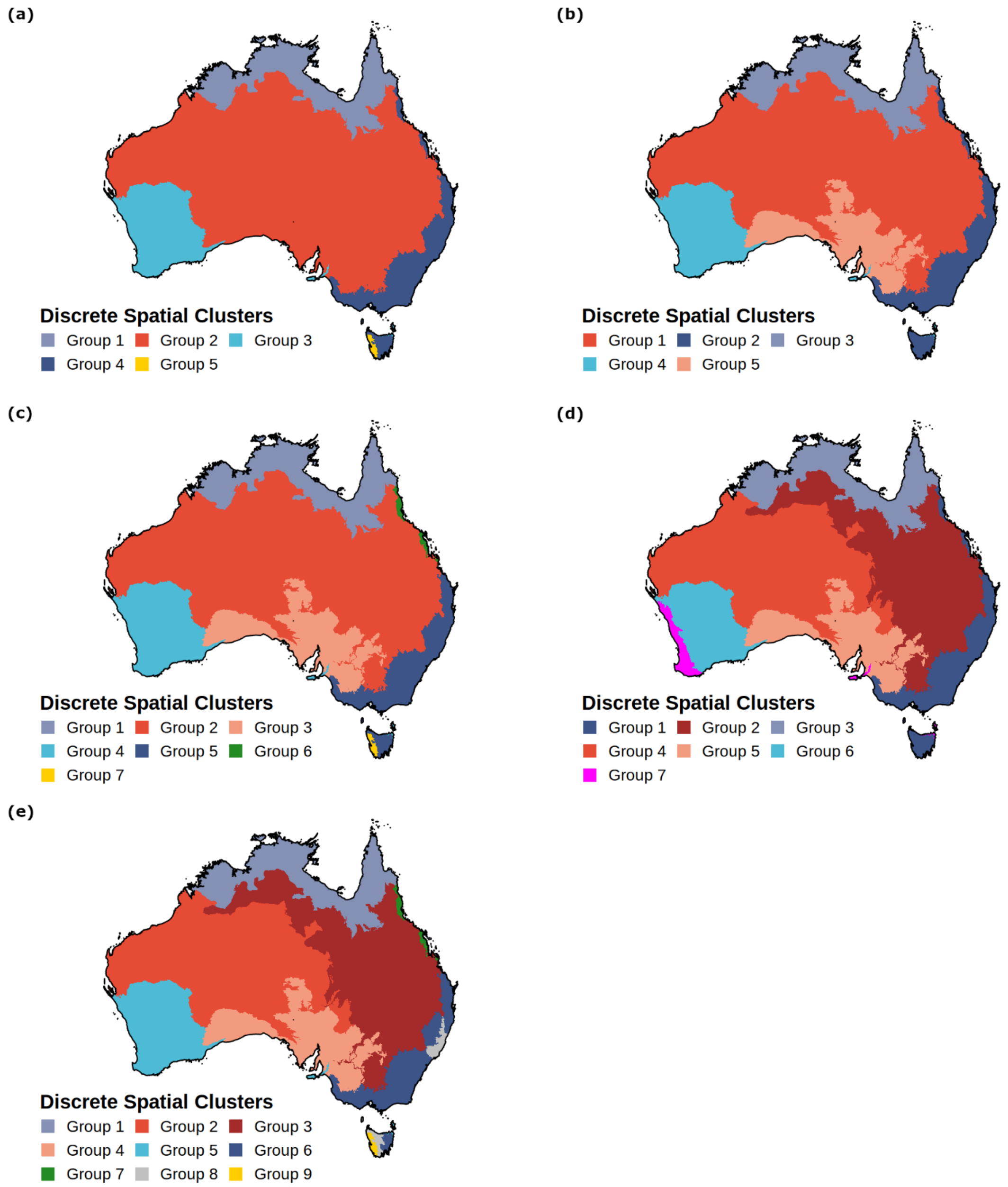


**FIGURE S5.5 Spatial nature of dissimilarity for the suite of optimal clusters of Australia's biophysical space.** Algorithm-driven aggregation of bioregions of the Interim Biogeographic Regionalization for Australia framework using the principal components of multiple factor analysis of a set of four geographic and environmental covariates (climate, lithology, soil, and vegetation). The resulting dendrogram was cut into (a and b) a pair of five and (c and d) another pair of seven groups using (a and c) the 'static' and (b and d) 'dynamic' tree-cutting techniques, and into (e) nine groups defined by the 'L' technique as the model that most accurately fitted the dataset while also minimizing complexity. Group 5, 7 and 9 in panels a, c, and e, respecitvely, include only one bioregion, thereby they were not considered in the identification of best SAR models as outlined in the suppementary material (Appendix S4). Color codes reflect the general geographic distribution of groups of bioregions in the Australian landscpe, although—where possible—colors match those used to differentiate WWF's biomes (Fig. S3.4 a).

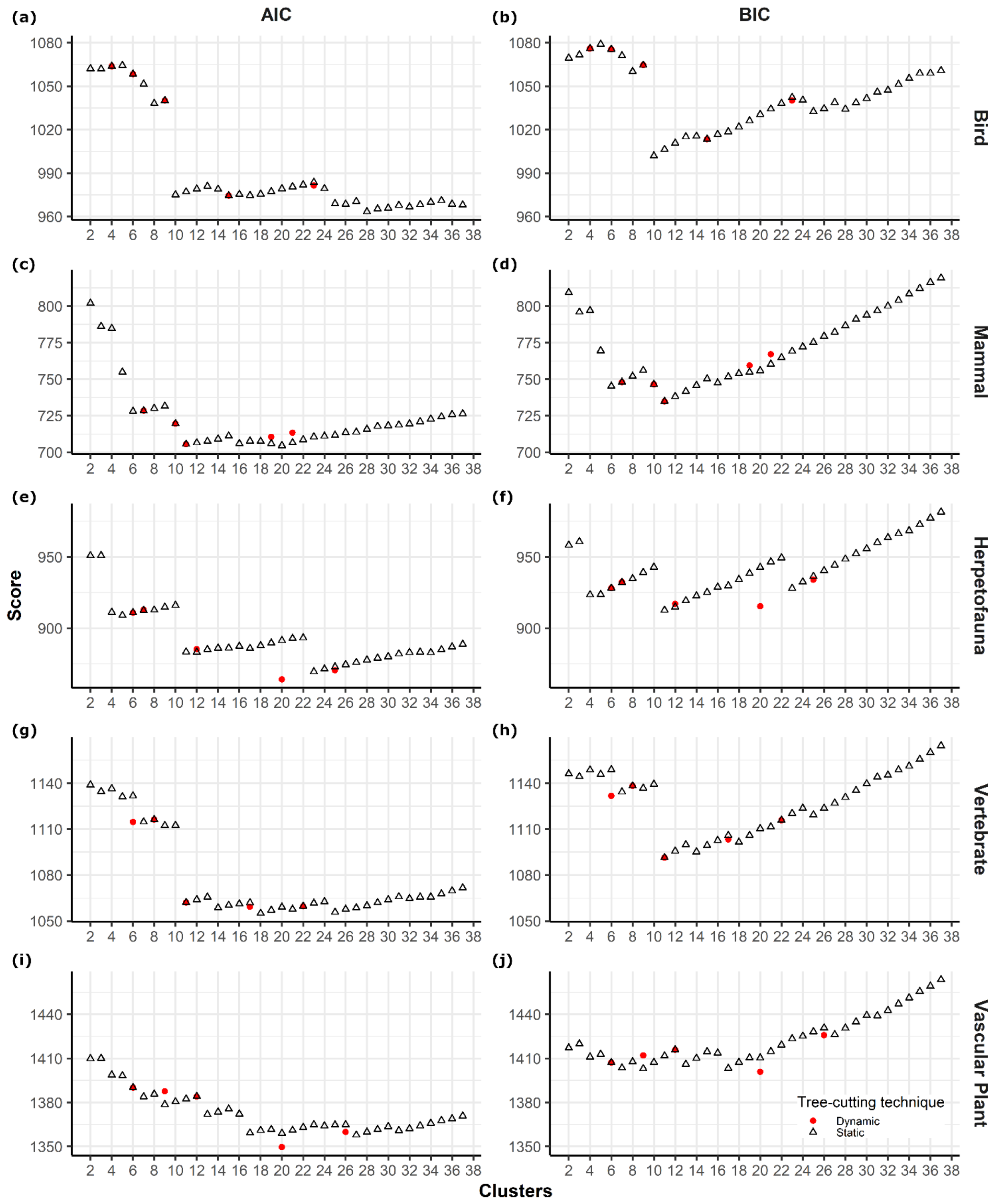


**FIGURE S6.6 Evaluation plots for alpha diversity.** Akaike's Information Criteria (AIC) and Bayesian Information Criteria (BIC) of the within- and across variance (ANOVA) in species richness for (a and b) birds, (c and d) mammals, (e and f) herpetofauna, (g and h) vertebrate, and (i and j) vascular plants against the number of clusters defined by the 'dynamic' and the 'static' tree-cutting techniques of the hierarchical cluster analysis of the compositional dissimilarity of the bioregions of the Interim Biogeographic Regionalization for Australia framework.

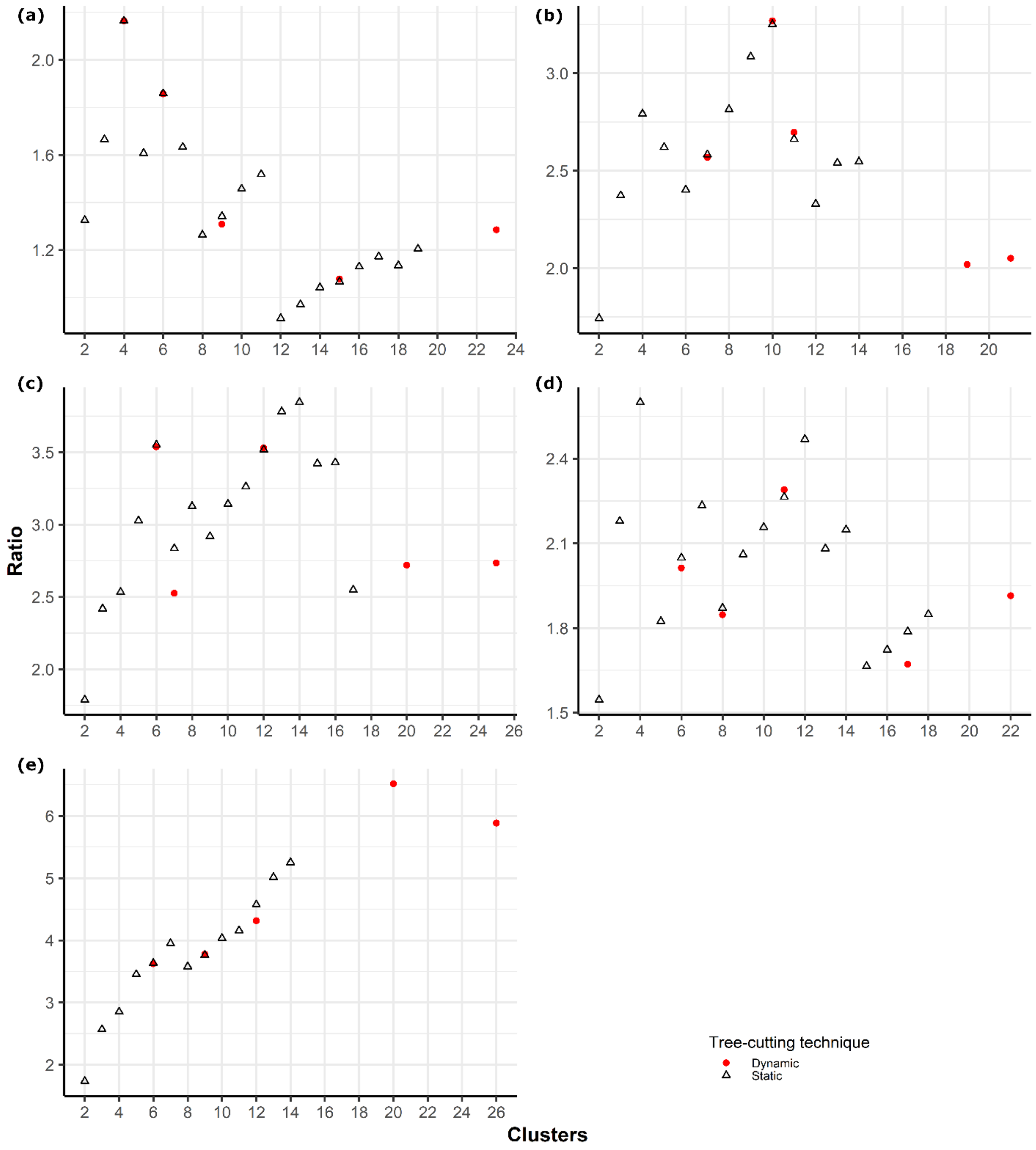


**FIGURE S7.7 Evaluation plots for beta diversity.** The ratio between the sum of multiple-site measures of compositional dissimilarity across groups of bioregions in the set of prominent clusters and the multiple-site measure of compositional dissimilarity across Australia's bioregions for (a) birds, (b) mammals, (c) herpetofauna, (d) vertebrate, and (e) vascular plants against the number of clusters defined by the 'dynamic' and the 'static' tree-cutting techniques of the hierarchical cluster analysis of the compositional dissimilarity of the bioregions of the Interim Biogeographic Regionalization for Australia framework with multiple bioregions in clusters. Only prominent clusters with all their groups including two or more bioregions were considered.

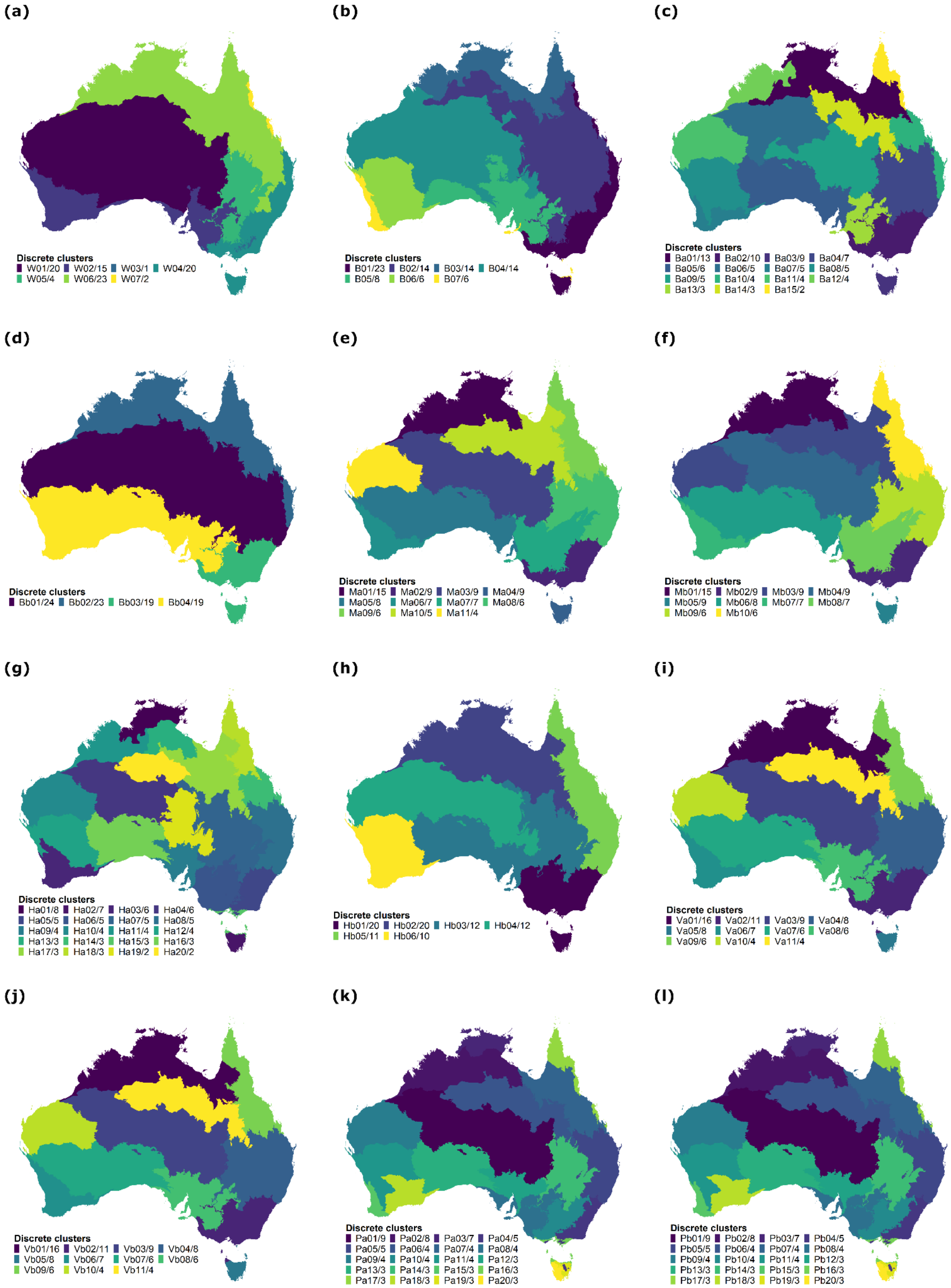


**See next page for figure caption**

**FIGURE S8.8 Spatial configuration of an expert-based and an algorithm-driven partition of Australia's bioregions.** Maps corresponding to (a) the World Wildlife Fund's expert-driven classification of Australia into seven biomes (Dinerstein et al., 2017), and (b) the optimal discrete-species clusters based on an algorithm-driven aggregation of the bioregions of the Interim Biogeographic Regionalization framework for a suite of geographic and environmental covariates (i.e., the optimal model/partition for the species-area relationship), or the variance in (c, e, g, i and k) alpha and (d, f, h, j and l) beta diversity based on the dissimilarity of species compostion for (c and d) bird, (e and f) mammal, (g and h) herpetofauna, (i and j) vertebrate, and (k and l) vascular plant species. Alphanumeric value identifies distinct clusters, and the number after slash corresponds to the number of bioregions in cluster.
